# Positively Selected Enhancer Elements Endow Tumor Cells with Metastatic Competence

**DOI:** 10.1101/155416

**Authors:** James J. Morrow, Ian Bayles, Alister PW Funnell, Tyler E. Miller, Alina Saiakhova, Michael M. Lizardo, Cynthia F. Bartels, Maaike Y. Kapteijn, Stevephen Hung, Arnulfo Mendoza, Daniel R. Chee, Jay T. Myers, Frederick Allen, Marco Gambarotti, Alberto Righi, Analisa DiFeo, Brian P. Rubin, Alex Y. Huang, Paul S. Meltzer, Lee J. Helman, Piero Picci, Henri Versteeg, John Stamatoyannopoulos, Chand Khanna, Peter C. Scacheri

## Abstract

Metastasis results from a complex set of traits acquired by tumor cells, distinct from those necessary for tumorigenesis. Here, we investigate the contribution of enhancer elements to the metastatic phenotype of osteosarcoma. Through epigenomic profiling, we identify substantial differences in enhancer activity between primary and metastatic tumors in human patients as well as near-isogenic pairs of high and low lung-metastatic osteosarcoma cells. We term these regions Metastatic Variant Enhancer Loci (Met-VELs). We demonstrate that these Met-VELs drive coordinated waves of gene expression during metastatic colonization of the lung. Met-VELs cluster non-randomly, indicating that activity of these enhancers and their associated gene targets are positively selected. As evidence of this causal association, osteosarcoma lung metastasis is inhibited by global interruptions of Met-VEL-associated gene expression via pharmacologic BET inhibition, by knockdown of AP-1 transcription factors that occupy Met-VELs, and by knockdown or functional inhibition of individual genes activated by Met-VELs, such as F3. We further show that genetic deletion of a single Met-VEL at the *F3* locus blocks metastatic cell outgrowth in the lung. These findings indicate that Met-VELs and the genes they regulate play a functional role in metastasis and may be suitable targets for anti-metastatic therapies.

## Introduction

More than 90% of all cancer deaths are the result of tumor metastasis^1^. The physical process of tumor cell dissemination and metastatic colonization of distant secondary sites has been well described^2^. Whole genome sequencing studies have elucidated the evolutionary phylogeny of metastatic dissemination^3, 4^, and gene expression studies have revealed many of the genes that mediate the progressive steps of metastasis and drive organ-specific colonization^5-7^. These studies suggest that adaptation of metastatic tumor cells to the microenvironments of their destination organs is accompanied by a shift in cell state. Whether the shift is driven by genetic or epigenetic factors, or a combination of both of these mechanisms is not yet clear.

During normal development, gene expression changes that accompany cell state transitions are driven by altered activity of gene enhancer elements^8-10^. Enhancers govern cell type-specific expression programs and are defined by signature chromatin features including H3K4me1, H3K27ac, and DNase hypersensitivity^11^. Enhancers appear to be important in tumorigenesis as well. Previous studies by our group have demonstrated that malignant transformation is accompanied by locus-specific gains and losses in enhancer activity across the epigenome. We previously termed these Variant Enhancer Loci (VELs)^12,13^. Others have shown that in many types of cancers, clusters of active enhancers called super-enhancers (SEs) mediate dysregulated expression of oncogenes^14,15^. Collectively, these studies suggest that aberrant enhancer activity is a key driver of tumor formation and maintenance. Further, these data show that enhancer profiling is a powerful approach for interrogating the transcriptional circuitry of malignant cell states as it aids in the identification of both cancer dependency genes and the transcription factors regulating their expression.

Altered transcriptional programs are known to play a role in metastatic tumor progression. In certain model systems, these transcriptional programs have been associated with metastatic colonization of specific secondary organs^5-7,16^. Recently, epigenetic changes have been associated with transcriptional changes during metastasis^17^. However, the contribution of gene enhancers to metastatic transcription is not well understood. Based on the knowledge that enhancers drive cell-state transitions during normal development and tumorigenesis, we hypothesized that enhancers may play a similar role in the transition of cancer cells from one developmentally distinct tissue to another during metastatic progression.

Osteosarcoma is the most common primary malignancy of the bone with peak incidence in children and adolescents. Clinical outcomes for patients have not improved for 30 years and there are currently no approved targeted anti-metastatic therapies for osteosarcoma in wide clinical use^18^. More than 75% of osteosarcoma metastases occur at the secondary site of the lung, which is the cause of the overwhelming majority of osteosarcoma related deaths^19^. In this study, we leverage the knowledge that gene enhancer activity is the cornerstone of cellular phenotypes and cell type specific gene expression^9,20^ to gain new insight into the regulatory mechanisms that allow metastatic osteosarcoma cells to overcome the critical barriers to colonization encountered as these cells engage the lung microenvironment. Our studies establish that enhancer elements endow tumor cells with metastatic capacity and that targeted inhibition of genes associated with enhancer alterations, or deletion of altered enhancers themselves is sufficient to block metastatic colonization and proliferation.

## Results

### The Metastatic Phenotype of Human Osteosarcoma is Associated with Variant Enhancer Loci

We mapped the locations of putative enhancer elements genome wide through ChIP-seq of the canonical enhancer-histone marks, H3K4me1 and H3K27ac in matched primary tumors and lung metastases from five osteosarcoma patients. We also performed H3K4me1 and H3K27ac ChIP-seq, and DNase-seq on a panel of five well-characterized ^21^ metastatic/non-metastatic human osteosarcoma cell line pairs representing three distinct mechanisms of metastatic derivation (Fig. 1a). Based on the previous finding that H3K4me1 broadly correlates with both poised and active enhancers^22,23^, we used this histone mark for our initial comparisons.

**Figure 1:**
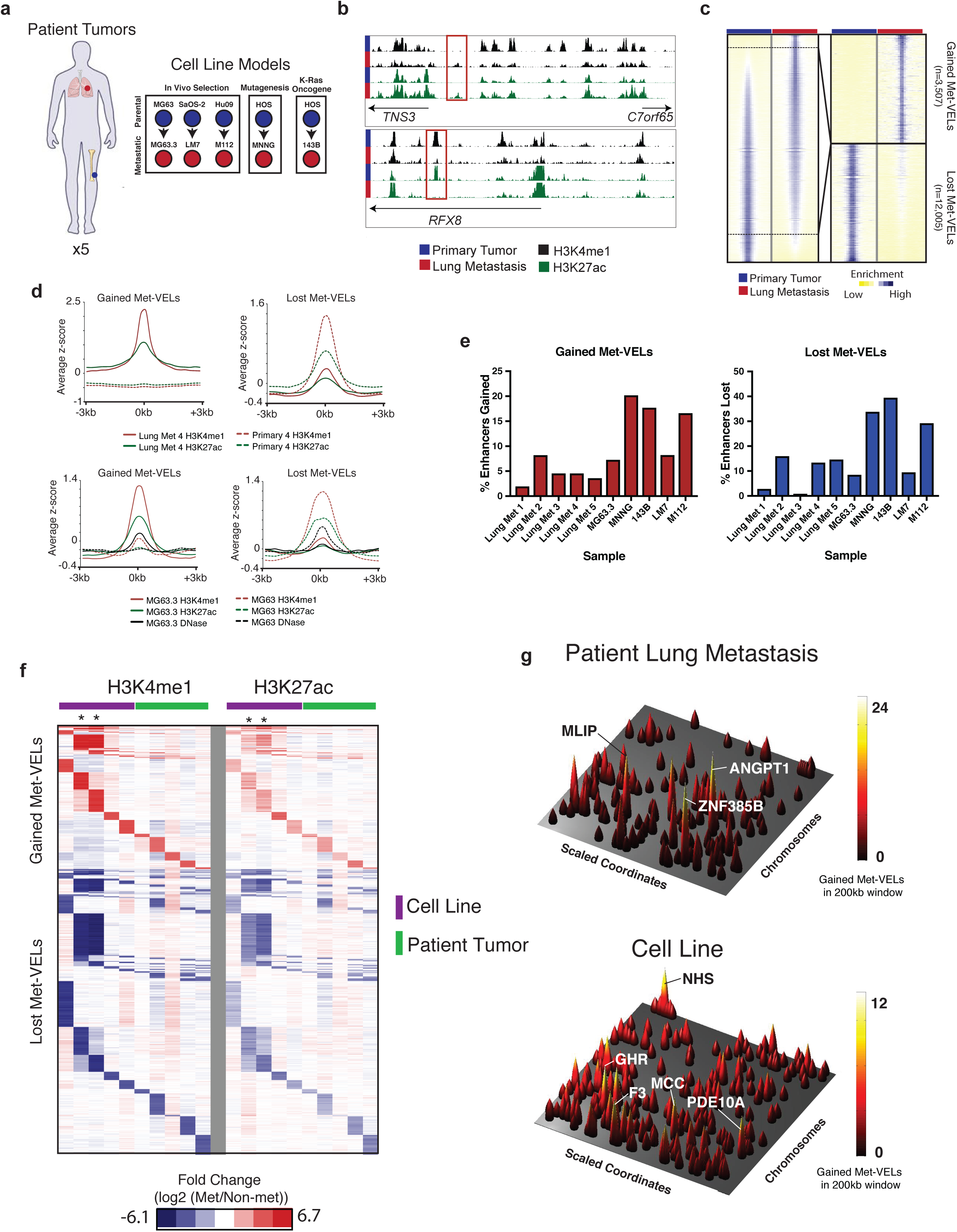
H3K4me1 ChIP-seq identifies metastatic variant enhancer loci (Met-VELs) and MetVEL clusters. **a,** Schematic representation of human tumor and metastatic human osteosarcoma cell line cohort. **b,** UCSC browser views of H3K4me1 profiles from MG63.3 (metastatic) and MG63 (parental) cell lines illustrating an example of gained (top) and lost (bottom) Met-VEL(s). Met-VELs are boxed in red. **c,** Heatmap showing H3K4me1 ChIP-seq signal +/-5kb from H3K4me1 peak midpoints for all putative enhancers in MG63.3/MG63 pair sorted by differences in signal. Sub-panel shows heatmap for gained and lost Met-VELs alone. **d,** Aggregate plots showing H3K4me1 ChIP-seq and H3K27ac ChIP-seq signal +/- 3kb from midpoints of gained (left) and lost (right) Met-VELs for a representative matched primary/lung metastatic human tumor pair (top) and MG63.3/MG63 cell line pair (bottom). DNase-seq signal +/- 3kb from Met-VEL mid-points is also shown for the MG63.3 and MG63 cell lines. **e.** Percentage of enhancers gained and lost in metastatic samples relative to primary tumors or non-metastatic cell lines. **f.** Heatmap of fold change normalized RPKM in metastatic samples vs. primary tumor or non-metastatic cell lines for aggregated list of all gained and lost Met-VELs across all samples. H3K4me1 signal shown on the left, H3K27ac signal shown on the right. The samples from left to right are as follows M112, 143B, MNNG, MG63.3, LM7, Lung Met 1, Lung Met 2, Lung Met 3, Lung Met 4, Lung Met 5. Asterisks indicate 143B and MNNG samples. **g.** Genome-wide gained Met-VEL landscape for human osteosarcoma metastatic tumor (Lung Met 4) and MG63.3 cell line. Rows represent scaled chromosomal coordinates. Peaks represent maximum gained Met-VEL counts in 200kb sliding windows. Predicted target genes for selected peaks are labeled.

In each comparison, we found thousands of regions where H3K4me1 signals showed at least a 3-fold difference in enrichment between conditions (Fig. 1b, 1c). These sites were either “gained” or “lost” in the metastatic cells relative to their matched controls. These metastasis-associated gains and losses of the H3K4me1 signal were reminiscent of those that we previously identified in the setting of primary tumor development through comparisons of primary colon tumors and normal colon tissue, known as VELs^12,13^. In distinction, we now and herein term the regions that show differential enrichment of H3K4me1 between metastatic samples and non-metastatic controls Metastatic Variant Enhancer Loci, or Met-VELs. Enhancers defined by differential enrichment of H3K4me1 generally showed concordant changes in H3K27ac ChIP-seq signals and DNaseseq signals (Fig. 1d and Extended Data Fig. 1), indicating robust commissioning and decommissioning of active enhancer elements at these loci. Across all samples, we found that on average 9.3% of all enhancers in a given metastatic cell line or tumor were gained relative to controls while 16.4% of enhancers present in non-metastatic cell lines or primary tumors were lost (Fig. 1e).

We next assessed the degree of Met-VEL heterogeneity across the cohort (Fig. 1F). We found that Met-VELs were more concordant between two metastatic cell lines (MNNG and 143B, labeled with asterisks) derived from a single parental cell line (HOS) than those derived from distinct parental cell populations (40.8% versus 0.2-9.3%, P < 0.001), suggesting that the specific enhancer elements that undergo activation and silencing is non-random and may be predetermined by the genetic and/or epigenetic makeup of the parental cell line. Met-VELs were heterogeneous among the remaining samples, with 23-69.1% showing overlap with at least one other sample, and 4.2-18.4% showing overlap with 2 or more samples. No Met-VELs were common to all cell lines or tissue samples.

An initial survey of Met-VEL distributions revealed dense clusters at distinct regions across the epigenome often in the vicinity of individual genes (Extended Data Fig. 2a). This finding led us to hypothesize that enhancer activity in these regions was non-randomly acquired due to selective pressures incurred during the process of metastatic progression. We systematically tested this hypothesis and found numerous loci with Met-VEL counts significantly greater than expected by chance in all samples (clusters in exemplar pairs displayed in Fig. 1g and Extended Data Fig. 2c). Several of the genes associated with Met-VEL clusters in both primary human samples and in the cell lines have been previously implicated in tumor biology and/or progression. ANGPT1, associated with >20 gained Met-VELs in one of the primary tumor/metastatic pairs, is a TIE2 receptor agonist that plays a crucial role in angiogenesis and is currently being studied as a therapeutic target in malignancy^24^. Among the genes associated with gained Met-VEL clusters containing the highest gained Met-VEL counts in the MG63.3 cell line pair were growth hormone receptor (*GHR*), phosphodiesterase 10A (*PDE10A*), and tissue factor (*F3*), all of which have been previously implicated in tumor biology and/or progression. GHR is a key signaling node in the growth hormone pathway. Activation of this pathway has been associated with tumor growth and acquisition of metastatic traits in multiple types of cancer^25^ including osteosarcoma^26^. PDE10A is a phosphodiesterase isozyme that is typically limited to expression in brain and testes tissues. PDE10A has been associated with colorectal cancer cell growth^27^, but has not been previously implicated in metastasis or associated with osteosarcoma. F3 is a well-described activator of normal blood coagulation. In the setting of cancer, F3 plays tumor-cell-endogenous roles in promoting tumor growth and metastasis in multiple cancers, but the mechanism underlying its activation is not fully defined^28^. We performed chromatin conformation capture studies and found that the majority of constituents of the Met-VEL cluster downstream of *F3* physically contact the transcription start site indicating coordinated activity of these non-randomly acquired enhancers (Extended Data Fig. 2b). All metastatic/non-metastatic cell line and primary/metastatic tumor pairs in our panel showed evidence of nonrandom acquisition and loss of enhancer clusters (Extended Data Fig. 2d). On average across the cohort, 22% of all Met-VELs were found to reside in Met-VEL clusters (Extended Data Fig. 2e).

We next sought to determine whether Met-VELs and Met-VEL clusters were unique to osteosarcoma or if this same process may occur in other tumor types as well. To test this, we analyzed previously published H3K4me1 ChIP-seq data from a matched pair of colon cancer cell lines derived from a primary tumor and a paired liver metastasis of the same patient^12^ and performed H3K4me1 ChIP-seq on flash-frozen tissue from a primary endometrial tumor and paired lymph node metastasis. We identified gained and lost Met-VEL clusters in metastatic colon cancer cells, and gained Met-VEL clusters in metastatic endometrial cancer (Extended Data Fig. 2f, g). Thus this feature is not limited to osteosarcoma but is likely to occur during the natural progression of multiple human tumor types. Through comparative analyses we further demonstrate that 77.1-98.5% of the Met-VELs detected in the osteosarcoma samples were not detected as Met-VELs in either the colon or endometrial tumor samples.

### Metastatic Variant Enhancer Loci (Met-VELs) Dynamically Modulate Gene Expression as Tumor Cells Engage the Lung Microenvironment

In order to investigate the role of Met-VELs in modulating gene expression during metastasis, we took advantage of an *ex vivo* murine model of osteosarcoma lung metastasis previously developed by our group^29^. In this model, we seed GFP-expressing tumor cells to the lungs of immunocompromised mice by intravenous tail vein injection and culture lung sections *ex vivo*. This approach allows us to track metastatic outgrowth of GFP-labeled tumor cells as they interact with the lung microenvironment in real time and to assess dynamic changes in gene expression throughout the process of metastatic colonization by transcriptome profiling of GFP-positive cells isolated by FACS (Fig. 2a). We performed RNA-seq at both early (24-hours) and late (day 14) time points in three cell line pairs. In all cases, genes associated with gained Met-VELs and Met-VEL clusters were generally expressed at higher levels in metastatic cells within the lung microenvironment than in their corresponding non-metastatic parental cell lines while genes associated with lost Met-VELs and lost Met-VEL clusters were expressed at lower levels (Fig. 2b and Extended Data Fig. 3a, b). To investigate whether Met-VEL-associated gene sets represent a transcriptional program specifically modulated in the setting of metastasis, we compared expression across conditions. The degree of differential expression of Met-VEL associated genes in the parental vs. metastatic cells was greater in *ex vivo* lung culture than in standard *in vitro* culture conditions (Extended Data Fig. 4a, b) indicating that modulation of these transcriptional programs represents a cellular response to external cues from the lung microenvironment. Met-VEL associated gene sets showed little overlap (< 27%) with the most differentially expressed genes in each metastatic/non-metastatic cell line pair (Extended Data Fig. 4c, d).

**Figure 2:**
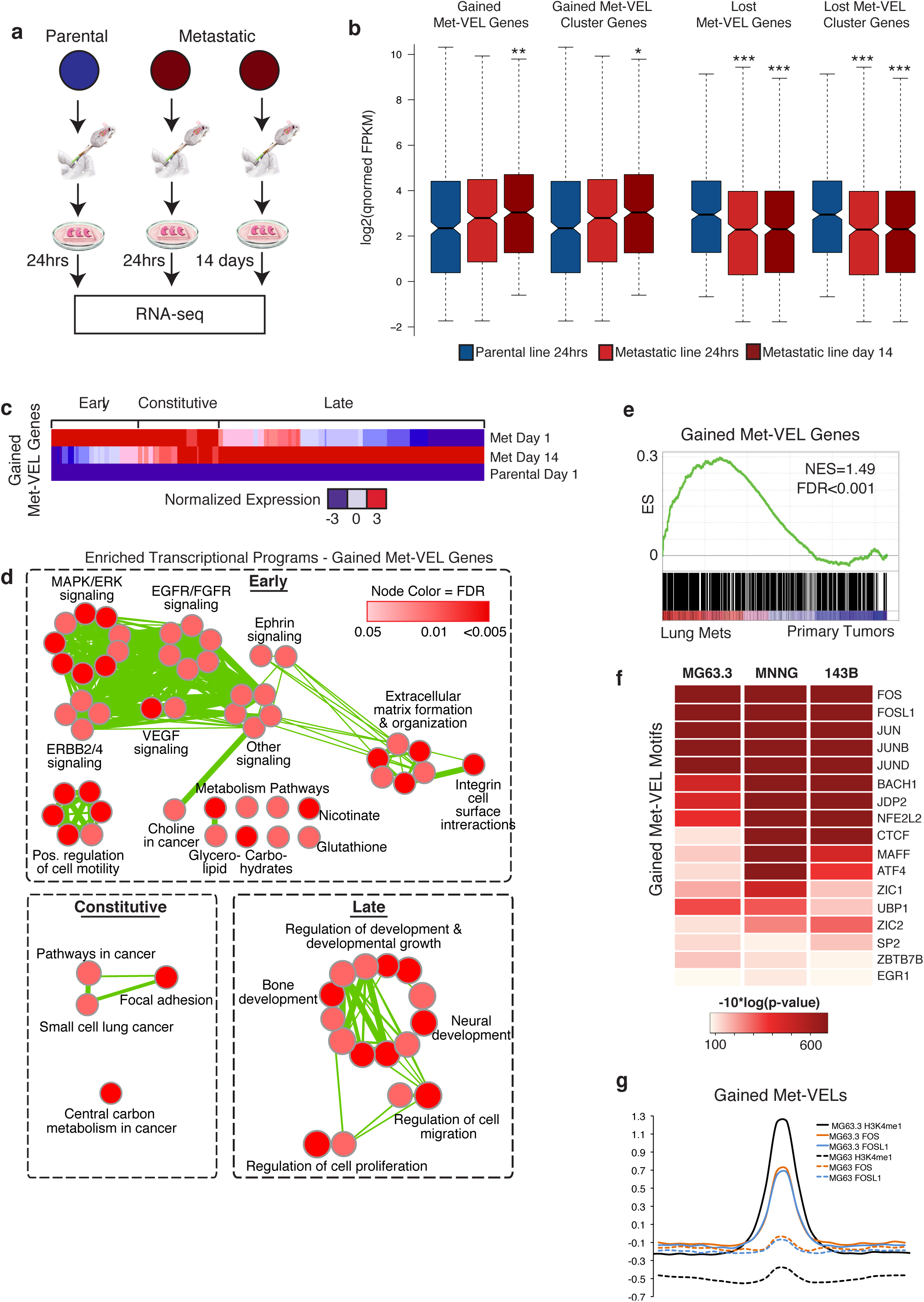
Met-VELs modulate gene expression during metastatic colonization of the lung. **a.** Schematic of experimental design for assessment of Met-VEL gene expression in parental and metastatic cell lines in *ex vivo* lung metastasis model. Image adapted from 29. **b.** Log2 quantile-normalized FPKM values for gained (left) and lost (right) Met-VEL and Met-VEL cluster genes in MG63/MG63.3 cell line pair. Asterisks indicate significant differences in FPKM distributions between parental and metastatic cell lines (* P<0.05; ** P<1 E-3; *** P<1 E-4). P-values calculated by Mann-Whitney Test. **c.** Heatmap of up-regulated gained Met-VEL genes in MG63/MG63.3 cell lines illustrating phasic expression pattern. **d.** Enriched Map representation of all Gene Ontology (GO) terms for three classes of gained MetVEL responder genes calculated by aggregating gene lists from all three cell line pairs. **e.** GSEA plot of up-regulated gained Met-VEL gene set compiled from three metastatic cell lines in human patient lung metastases versus primary tumors. **f.** Expressed transcription factors with enriched motifs in gained Met-VELs in three metastatic/parental cell line pairs and corresponding motif enrichment p-values. **g.** Aggregate plot of H3K4me1, FOS, and FOSL1 ChiP-seq signal at all gained Met-VELs in MG63.3 cell line.

We found that subsets of genes associated with gained Met-VELs became highly expressed within 24hrs of arrival of metastatic cells to the lung, others were only activated later during metastatic outgrowth, and a third subset were constitutively up-regulated (Fig. 2c and Extended Data Fig. 3c). We assessed these gene sets for functional enrichment, and the results indicate that Met-VELs coordinate phasic waves of gene expression associated with progressive stages of lung colonization (Fig. 2d). Early responder genes were enriched for functions related to intracellular signaling pathways, metabolism, and cell surface interactions. Late responder genes were enriched for functions related to development, migration, and proliferation. Constitutively overexpressed genes were enriched for canonical cancer pathways, focal adhesion, and carbon metabolism. Functional enrichments of lost Met-VEL genes included embryonic morphogenesis and extracellular matrix production (Extended Data Fig. 3e). We verified that gained Met-VEL genes upregulated in the *ex vivo* lung model are frequently elevated in osteosarcoma patient lung metastases relative to primary tumors (Fig. 2e) and many of the same gene sets are enriched in gained Met-VEL target genes in osteosarcoma lung metastases in patients (Extended Data Fig. 5), thereby verifying that the genes identified using the cell line models and the *ex vivo* approach are representative of those dysregulated in human patients.

To identify transcription factors (TFs) that may drive Met-VEL expression programs, we performed motif enrichment analysis and identified a number of commonly expressed TFs with enriched motifs in gained and lost Met-VELs across all three pairs analyzed. The most highly enriched motifs include many members of the AP-1 complex (JUN, JUNB, JUND, FOS, and FOSL1) (Fig. 2f and Extended Data Fig. 3d) which has been previously shown to play a key role in osteosarcoma metastasis^30^. Intriguingly, we found that AP-1 motifs were enriched at both gained and lost Met-VELs. This finding suggests that Met-VELs likely alter the transcriptional programs mediated by AP-1 during osteosarcoma metastasis via a shift in target gene regulation within metastatic cells. We verified that FOS and FOSL1 are bound at gained Met-VELs by ChIP-seq for these transcription factors (Fig. 2g).

### Met-VEL Associated Gene Expression is Required for Metastatic Colonization

We used the BET-inhibitor JQ1 as a chemical tool to investigate the functional relevance of Met-VELs. JQ1 has previously been shown to inhibit osteosarcoma primary tumor formation through its effects on both tumor cells and bone cells within the tumor microenvironment^31^. We found that JQ1 showed potent anti-proliferative effects on metastatic tumor cells growing in the lung microenvironment without affecting the surrounding normal lung tissue (Extended Data Fig. 6a-c). The anti-proliferative effects were associated with selective suppression of gained Met-VEL target genes that are normally up-regulated in metastatic cells in response to cues from the lung microenvironment (Extended Data Fig. 6d, e). Intriguingly, JQ1 more potently suppressed gained Met-VEL genes than genes associated with super enhancers (Extended Data Fig. 6f) that have previously been linked to JQ1’s anti-tumor properties in other tumor models^15,32,33^.

The results of the JQ1 studies raise the possibility that metastatic outgrowth is dependent on appropriate activation of genes associated with gained Met-VELs. To test this hypothesis, we conducted a functional *in vivo* RNAi assay. We reasoned that if single genes and the Met-VELs driving their expression are the basis for selectable traits of the metastatic phenotype, then lack of expression of these genes should be deleterious to cells over the course of metastatic progression. We constructed a custom shRNA library for this assay targeting 33 genes. This gene list included 20 genes associated with gained Met-VELs or Met-VEL clusters, 11 TFs with motifs enriched at Met-VELs, and 2 genes of interest from other ongoing studies. We cloned the shRNA library into a Tet-ON lentiviral construct (LTREPIR, Fig. 3a) modified from a similar construct previously published^34^. Using a DsRed fluorescent reporter of shRNA induction, we show that this construct was robustly induced upon exposure to doxycycline and not leaky in the absence of doxycycline (Extended Data Fig. 7). We conducted parallel screens *in vivo* and *in vitro* to allow us to distinguish genes that specifically inhibit metastatic outgrowth in an *in vivo* microenvironment (i.e. metastasis dependency genes) from genes whose inhibition reduces cellular growth independent of context (Fig. 3b). In the *in vivo* screen, transduced cells were delivered via tail vein injection into mice pre-treated with doxycycline. Mice were maintained on doxycycline throughout the 21-day course of the experiment. In the parallel *in vitro* screen, transduced cells were treated in culture with doxycycline for 21 days. At the conclusion of the experiment, induced cells actively expressing an shRNA (DsRed+/GFP+) were sorted from mouse lungs or *in vitro* culture by FACS. DNA was isolated from these cells, along with uninduced cells from the initial population (input), and shRNAs were amplified, sequenced, and aligned to the reference shRNA sequences of the library to determine normalized representation of each shRNA.

**Figure 3:**
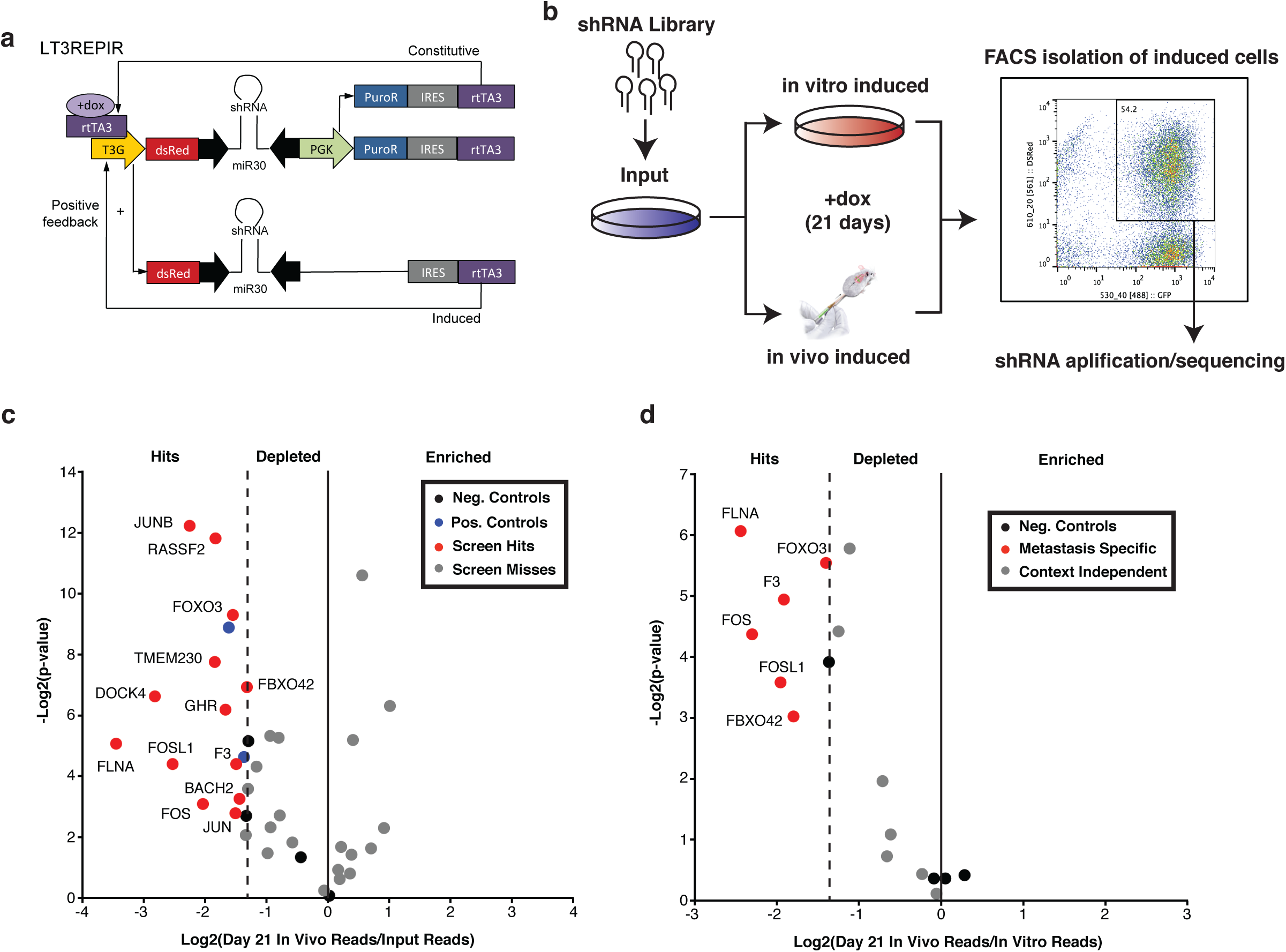
*In Vivo* high-throughput RNAi functional assay of candidate metastasis dependency genes. **a.** Schematic of doxycycline-inducible LT3REPIR shRNA construct. Modified from 34. **b.** Schematic of experimental design for *in vivo* high-throughput functional assay of candidate metastasis dependency genes. **c.** Volcano plot of relative abundance of shRNAs targeting 33 genes in GFP+/DsRed+ sorted osteosarcoma cells from doxycycline-treated mice (N=5 per replicate x 3 replicates) versus input cell population. 2^nd^ most depleted shRNAs for each gene are plotted as well as negative and positive shRNA controls. Negative controls contained groups of 2-4 shRNAs. **d.** Volcano plot of relative abundance of shRNAs targeting 13 genes meeting initial hit criteria (Fig. 3c) in GFP+/DsRed+ sorted osteosarcoma cells from doxycycline-treated mice (N=5 per replicate x 3 replicates) versus GFP+/DsRed+ sorted osteosarcoma cells treated with doxycycline *in vitro*. 2nd most depleted shRNAs for each gene are plotted as well as negative controls groups of 2-4 shRNAs.

We defined metastasis dependency genes as those whose knockdown inhibited *in vivo* metastasis significantly more than *in vitro* growth. First, shRNA representations in metastatic tumor cells were compared to input representations to identify shRNAs depleted from the population of cells during metastatic outgrowth in the lung (Fig. 3c). Genes that were targets of at least two non-overlapping shRNAs (the pool contained 3-4 shRNAs per gene) that inhibited metastatic outgrowth to a greater degree than all negative controls were defined as initial hits. Because we used a second filter for these hits in this screen and planned further functional validation experiments, we intentionally chose a relatively inclusive threshold for initial hit calling. We found that 13 of 33 genes (39%) included in our screen met this criterion. To determine if depletion of cells expressing these shRNAs was specific to metastasis, we compared the relative representations of hits in the *in vivo* induced population of cells to *in vitro* induced controls (Fig. 3d). Genes whose shRNAs were significantly more depleted *in vivo* than *in vitro* were considered metastasis dependency genes. 6 of the 13 initial hits met this criterion (Extended Data Table 1). Metastasis dependency genes included four genes associated with gained Met-VEL clusters (*F3*, *FBXO42*, *FLNA*, and *FOXO3*) as well as two AP-1 complex TFs whose motifs are enriched in Met-VELs and were shown to bind at these enhancers (*FOS* and *FOSL1*). These results indicate that metastatic colonization of the lung by osteosarcoma cells is dependent on expression of a subset of individual genes associated with gained Met-VEL clusters as well as AP-1 complex TFs likely to regulate Met-VEL transcriptional programs.

Among the Met-VEL genes, Tissue Factor (*F3*) emerged as a top candidate driver of metastasis in osteosarcoma. In the MG63.3 cell line, *F3* was associated with a gained Met-VEL cluster containing the second highest gained Met-VEL count of the entire data set (Fig. 1g), suggesting this locus was under particularly strong positive selection during metastatic derivation. Gained Met-VELs in the *F3* cluster also showed higher levels of H3K27ac and DNase accessibility in MG63.3 cells compared to the parental MG63 cell line and chromatin conformation capture studies confirmed that these enhancers physically contact the transcription start site of *F3* (Extended Data Fig. 2b). We also found that *F3* was more highly expressed in MG63.3 cells during metastatic outgrowth than in the parental MG63 cells (Extended Data Fig. 8a). In addition, two other metastatic cell lines, MNNG and 143B, showed active enhancer signals at the *F3* locus, similar to MG63.3 (Extended Data Fig. 8c) and expressed *F3* at higher levels during metastatic outgrowth than their parental cell line (Extended Data Fig. 8a). To verify that elevated *F3* transcript levels were recapitulated at the protein level and also not an artifact of *ex vivo* culture, we performed immunofluorescence analysis of lung metastases from a fully *in vivo* model of metastasis and confirmed that metastatic osteosarcoma cells expressed higher levels of F3 protein than non-metastatic cells (Fig. 4a, b). Quantification of *F3* levels directly in human osteosarcoma patient samples showed that *F3* was elevated in lung metastases relative to primary tumors (Extended Data Fig. 8b). Further, gained Met-VELs at the *F3* locus were detected in 3 of the 5 patient lung metastases (Extended Data Fig 9). Using a tissue microarray, we confirmed that F3 protein was highly expressed in lung metastases from human osteosarcoma patients. F3 was expressed in >50% of tumor cells in 18/18 lung metastases and F3 showed strongly positive staining in 17/18 samples (Fig. 4c, d).

**Figure 4:**
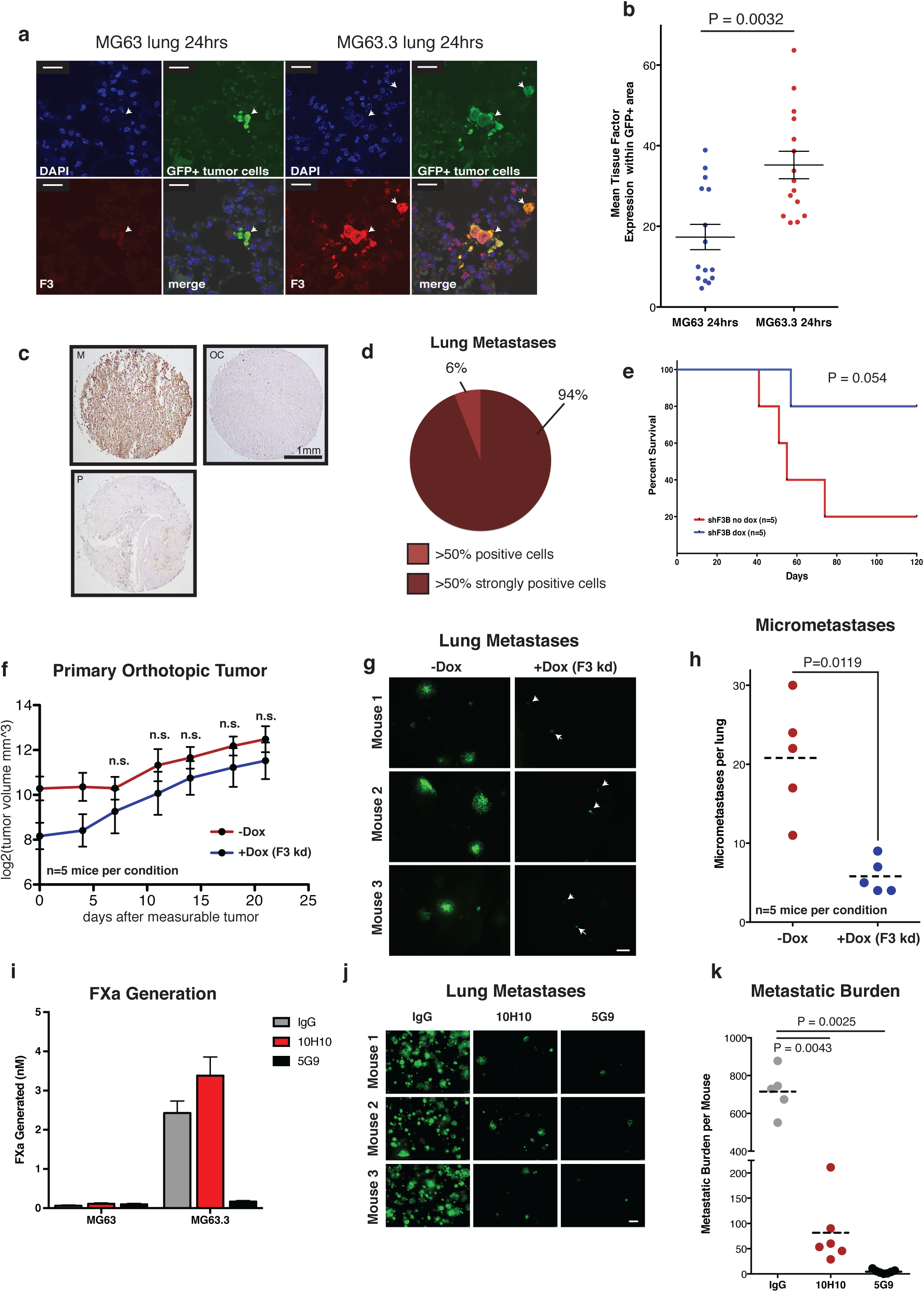
Tissue Factor (F3) mediates lung metastasis of osteosarcoma. **a.** *In Vivo* 2.5x images of GFP+ MG63 (parental) and MG63.3 (metastatic) cells in the lung 24hrs following tail-vein injection of 1x10^6^ cells. Sections stained for GFP, tissue factor (F3, red), and DAPI. Arrowheads indicate individual tumor cells within lung. Scale bars = 20μm. **b.** Quantification of mean red pixel intensity within GFP+ (tumor) area in MG63 (parental) and MG63.3 (metastatic) cells 24hrs after tail-vein injection (N=15 images per condition). **c.** Representative images of immunohistochemical staining of F3 in human osteosarcoma lung metastases (M), primary tumor (P), and omission control (OC). Tissue microarray contained 18 scoreable lung metastases of similar quality to those displayed. **d.** Percentage of lung metastases with various levels of F3 positivity. **e.** Kaplan-Meier plot of untreated (red) and doxycycline-treated (blue) mice tail-vein injected with 5x10^4^ GFP+ MG63.3 cells transduced with shF3B construct (N=5 mice per condition). P-value calculated by Gehan-Breslow-Wilcoxon Test. **f.** Primary tumor growth in untreated (red) and doxycycline-treated (blue) mice receiving orthotopic injection of 8x10^5^ GFP+ MG63.3 cells transduced with shF3B construct. Values represent averages +/-SEM (N=5 mice per condition). P-values calculated using student’s t-test. **g.** Representative 2.5x images of *in vivo* metastatic lesions in lungs 21 days after measureable tumor formation in untreated (left) and doxycycline-treated (right) mice receiving orthotopic injection of 8x10^5^ GFP+ MG63.3 cells transduced with shF3B construct. Arrowheads indicate individual tumor cells within lung. Scale bar = 500μm. **h.** Quantification of lung metastatic burden 21 days after measureable tumor formation in untreated (red) and doxycycline-treated (blue) mice receiving orthotopic injection of 8x10^5^ GFP+ MG63.3 cells transduced with shF3B construct (N=5 mice per condition, 5 images per mouse). P-values calculated using Mann-Whitney test. **i.** Amount of activated factor X (FXa) formed in *in vitro* assay by MG63 (left) and MG63.3 (right) cells treated with 25μg/mL IgG control, 10H10, or 5G9 antibodies for 20 minutes prior to the addition of FVIIa and FX to a final concentration of 100nM. FXa formation assessed 30 minutes after adding FX. P-values calculated using student’s t test with Welch’s correction. **j.** Representative 2.5x images of *in vivo* metastatic lesions in lungs of mice 14 days after tail vein injection of 5x10^5^ MG63.3 cells with 500μg of IgG, 5G9, or 10H10 antibodies. Scale bar = 500μm. **k.** Quantification of lung metastatic burden 14 days after tail vein injection of 5x10^5^ MG63.3 cells with 500μg of IgG, 5G9, or 10H10 antibodies (N= at least 5 mice per condition, 5 images per mouse). P-values calculated using Mann-Whitney test.

Having validated that F3 is dysregulated across multiple experimental model systems and shown that it is ubiquitously expressed at high levels in patient metastases, we sought to further test the functional contribution of F3 expression to the metastatic phenotype. We cloned two shRNAs targeting *F3* that were not included in the RNAi assay into the doxycycline inducible LTREPIR construct. We used an inducible vector for these experiments to ensure that alterations in metastatic competence could not be attributed to selection of transduced cell populations and used uninduced controls for all experiments. Relative to uninduced cells, *F3* expression was reduced 44-63% 40hrs after induction of each shRNA (Extended Data Fig. 10). F3 knockdown with these shRNAs did not affect the *in vitro* growth rate of metastatic MG63.3 or MNNG cells (Extended Data Fig. 11a), but significantly reduced metastatic outgrowth of these cells in *ex vivo* lung culture (Extended Data Fig. 11b, c), supporting the metastasis-specific role for F3 in this setting. Indeed, F3 knockdown also significantly reduced metastatic outgrowth of osteosarcoma cells *in vivo* (Extended Data Fig. 11d, e) and substantially prolonged survival of mice injected with metastatic osteosarcoma cells (Fig. 4e). To further test whether F3 knockdown reduces *in vivo* growth of metastatic cells generally or if this effect is specific to metastatic outgrowth of cells in the lung, we completed a spontaneous metastasis experiment using an orthotopic injection model. We found that F3 knockdown did not reduce primary tumor development or growth (Fig. 4f), but significantly inhibited metastasis, reducing average metastatic burden by 3.6 fold (Fig. 4g, h). While extensive GFP+ metastatic lesions were observed in control mice, lungs of mice in the F3 knockdown group were virtually devoid of metastatic lesions with only rare single GFP+ cells observable in most cases (Extended Data Fig. 12).

F3 is known to both induce blood coagulation by mediating the generation of the active form of factor X (FXa) in the coagulation cascade and to promote cell survival and proliferation upon binding to activated factor VII via intracellular signaling mechanisms^28^. To determine the relative contributions of each of these functions to the pro-metastatic role of F3 we took advantage of monoclonal antibodies generated to inhibit each of these functions independent of the other^35^. As expected, we found that MG63.3 cells produce more FXa in *in vitro* assays than MG63 cells (Fig. 4i). We confirmed that anti-coagulant Mab-5G9 robustly inhibited this activity while antibody Mab-10H10, designed to prevent intracellular signaling, did not. We next tested the anti-metastatic effects of these antibodies *in vivo*. We co-injected MG63.3 cells with each of these antibodies or IgG control into the tail veins of mice and found that both inhibited metastasis, with Mab-5G9 showing a more pronounced effect (Fig. 4j,k). These results indicate that both the intracellular signaling and pro-coagulant functions of F3 contribute to metastatic progression, but that F3’s pro-coagulant activity is especially critical to metastatic success. Collectively, these results suggest that Met-VELs regulate expression of genes, such as F3, with critical functions during metastatic progression.

In order to directly test the role of Met-VELs in mediating the metastatic phenotype we employed transcription activator-like effector nuclease (TALEN) genome editing to excise one of the gained Met-VELs predicted to regulate *F3* in the metastatic cells. We targeted a Met-VEL located in a particularly robust DHS site containing high levels of both H3K4me1 and H3K27ac (Fig. 5a). This site also showed high ChIP-seq enrichment of both FOS and FOSL1 AP-1 complex members. We generated a cell clone with homozygous deletion of this Met-VEL, verified by Sanger sequencing (Fig 5a). Edited and unedited control cells were then seeded to mouse lungs via tail vein injection and the growth of the cells was monitored in the lungs using the *ex vivo* metastasis assay. Quantification of F3 levels 24-hours post-injection showed that F3 expression was reduced by 34% in the edited cells relative to unedited control cells (Fig. 5b, c). By day 5, lungs seeded with the F3 Met-VEL-edited cells were nearly entirely devoid of tumor cells, while extensive GFP+ metastatic lesions were observed in lungs seeded with the unedited cells (Fig. 5d). Quantification showed that deletion of this gained Met-VEL decreased metastatic burden by 78% (Fig. 5e).

**Figure 5:**
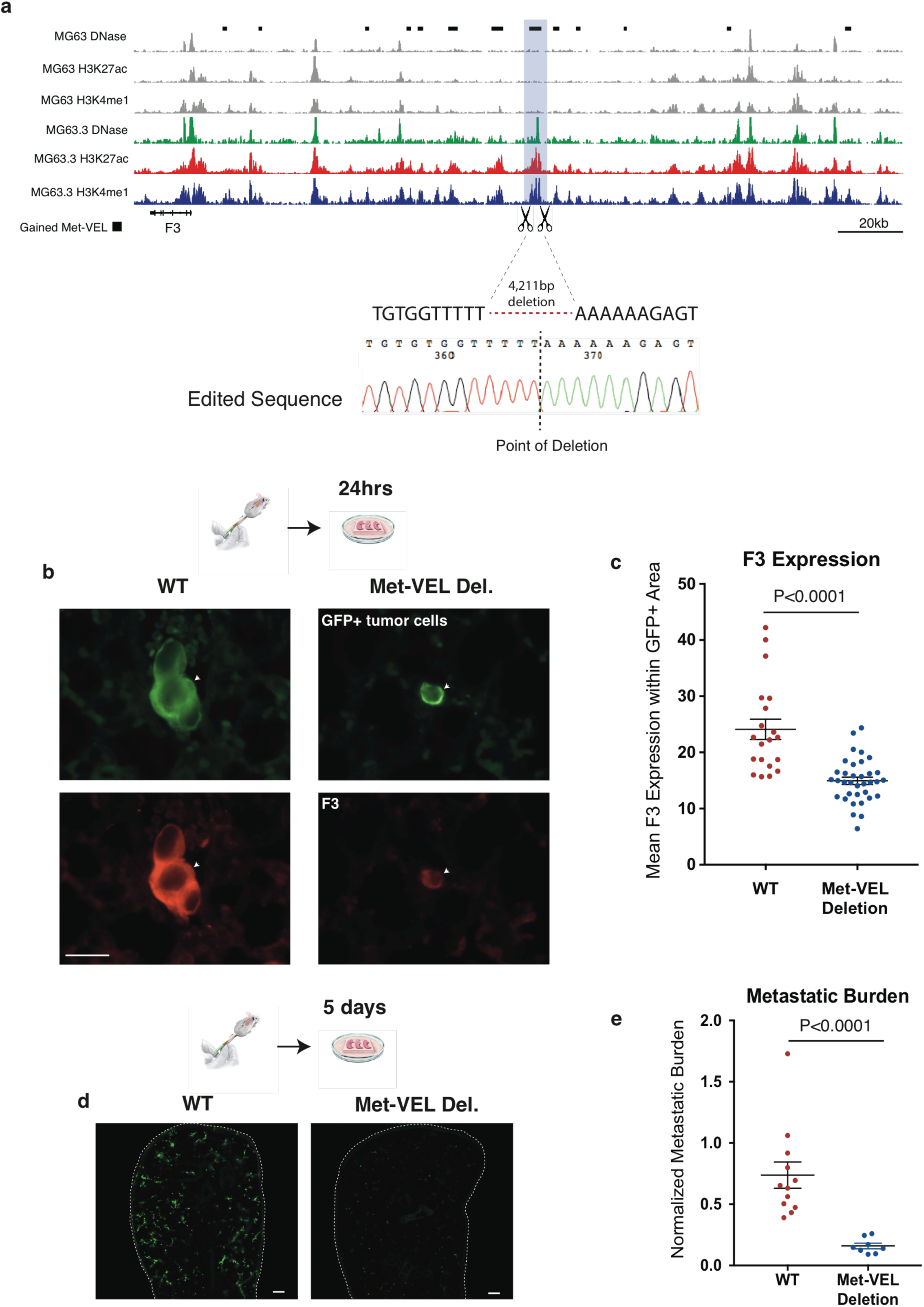
Deletion of single gained Met-VEL blunts F3 expression and mitigates lung metastasis of osteosarcoma cells. **a.** IGV browser view of region targeted for deletion with TALENs. Schematic shows strategy for 4,211 bp deletion. Sanger sequencing shows resulting clonal homozygous deletion. **b.** Representative 40x images of WT and Met-VEL deleted MG63.3 cells in the lung 24hrs after initiation of an ex vivo lung metastasis experiment. Scale bar = 50μm. **c.** Quantification of mean red pixel intensity (F3 expression) within GFP+ (tumor) area for WT and Met-VEL deleted MG63.3 cells 24hrs after initiation of *ex vivo* lung metastasis assay (N= at least 20 images per condition). P-value calculated by Mann-Whitney Test. **d.** Representative 2.5x images of WT (left) and Met-VEL deleted (right) lung sections at day 5. Lung sections outlined with dashed white line. Scale bar = 500μm. **e.** Quantification of metastatic burden at day 5 of *ex vivo* lung culture for GFP+ WT and Met-VEL deleted MG63.3 cells. Bars represent mean +/-SEM from at least 8 sections per condition (4 sections per mouse x 2-3 mice) normalized to the same section at day 0. P-Value calculated by Mann-Whitney Test.

## Discussion

While many of the genes responsible for metastatic progression have been identified across tumor types, the underlying mechanisms regulating expression of these genes are not well defined. Our studies demonstrate that altered enhancer activity is a fundamental mechanism by which tumor cells regulate gene expression during the dynamic process of metastasis, and thereby acquire metastatic traits. Through epigenomic profiling experiments, we identify enhancers that distinguish human osteosarcoma lung metastases from matched primary tumors and verify that these differences are also present in near-isogenic metastatic/non-metastatic human osteosarcoma cell lines. Such enhancer differences were also observed in colorectal and endometrial cancer, suggesting this is a general feature of metastatic progression. Subsets of these enhancer changes occur in non-random clusters indicating that they were positively selected during the process of metastatic derivation. These results demonstrate that the metastatic phenotype is accompanied by a shift in the enhancer epigenome, similar to the enhancer shifts that occur as cells transition through successive stages of embryonic development^8-10^, or during conversion of a normal cell to the malignant state^12,14,15^. The findings suggest that the evolutionary selective forces encountered by tumor cells during metastasis act to shape the enhancer landscape of metastatically successful cancer cell populations. The result of this selection is a population of cells possessing all of the traits necessary to overcome the barriers to metastatic colonization at distant tissues. Indeed, we show that many genes previously associated with metastasis become dysregulated through alterations in enhancer activity.

We provide multiple lines of evidence that acquired enhancer changes in metastatic osteosarcoma cells are functional and relevant to the metastatic phenotype in experimental models and human tissues. First, using human xenograft models of lung metastasis, we show that Met-VEL genes are dynamically regulated as metastatic cells engage the lung microenvironment and proliferate. Interestingly, Met-VELs predict “phasic” genes that become transcriptionally active at both early and late stages of metastatic outgrowth in the lung. Second, we demonstrate that metastatic cell outgrowth in the lung can be mitigated with a BET-inhibitor, and that this effect is associated with selective suppression of genes that are normally activated by Met-VELs in the lung. Third, through *in vivo* functional RNAi-based assays, we demonstrate that the metastatic capacity of the osteosarcoma cells can be diminished by targeted inhibition of individual Met-VEL genes and associated AP1-family transcription factors that likely regulate Met-VEL transcriptional programs. Using a fully *in vivo* spontaneous model of metastasis, we further verify that one such Met-VEL gene, Tissue Factor (*F3*), is a clinically relevant, *bone fide* metastasis gene essential for metastatic colonization with no apparent advantage to growth of the primary tumor. Further, interrupting the signaling and pro-coagulant functions of F3 was sufficient to inhibit metastasis, shedding light on the biological role of this gene in the metastatic progression of osteosarcoma. Our genomic Met-VEL deletion experiments demonstrate that the loss of function of a single gained enhancer is sufficient to impair metastatic colonization and subsequent outgrowth in mice, indicating that enhancer activation contributed to acquisition of the metastatic phenotype of these cells. Collectively the findings indicate that altered enhancer activity is a driver of gene expression that is critical for tumor cells to overcome the barriers of distal tissue colonization during metastasis.

It is well established that primary tumor formation is driven by a combination of genetic and epigenetic events^36^. With respect to metastasis, studies have shown that primary and matched metastatic tumors are broadly similar at the genetic level with no recurrent mutations identified in metastases that were not present in the primary tumor^3,4,37-42^. These studies suggest that primary tumors are likely already genetically equipped with the ability to metastasize. Further this implies that epigenetic processes may mediate the shift in cell state that accompanies metastatic progression, as proposed by others^17,43-46^. Consistent with this epigenetic hypothesis, we show that metastasis is accompanied by a shift in epigenetic state at enhancer elements. While our findings are not mutually exclusive with genetic theories of metastatic progression, we find that positive selection of enhancer activity is a fundamental component of the metastatic phenotype.

## Methods

### Cell Culture

Human osteosarcoma cell lines HOS, SaOS, MG63, MNNG, and 143B were purchased from ATCC. Hu09 and M112 cell lines were obtained from Dr. Jun Yokota (National Cancer Center Research Institute, Tokyo, Japan). LM7 was obtained from Dr. Eugenie S. Kleinerman (The University of Texas: MD Anderson Cancer Center, Houston, TX). MG63.3 cells were derived from MG63.2 (obtained from Dr. Hue Luu, University of Chicago, Chicago, IL) by metastatic selection in mice as previously described^47^.

The metastatic properties of these clonally related parental and metastatic cell lines have been thoroughly characterized in multiple murine models of metastasis^21^. A sample size of 5 cell line pairs was chosen to capture the spectrum of methods of metastatic derivation and to sufficiently power the study for comparative analyses based on similar studies completed by our lab in the past.

All the cells were cultured in DMEM (Life Technologies) medium supplemented with 10% fetal bovine serum (FBS) and glutamine except for the Hu09 and M112 lines that were cultured in RPMI 1460 (Life Technologies) with 10% FBS and glutamine.

The purity and authenticity of all lines used in these studies has been independently confirmed by short tandem repeat (STR) profiling performed by the International Cell Line Authentication Committee. Mycoplasma testing was routinely performed with Mycoalert Mycoplasma detection kit (Lonza).

### Mouse Studies

All animals were housed and handled in accordance with protocols approved by the CWRU IACUC or the NCI IACUC depending on location of performed studies. The number of animals included in each of the described studies was based on extensive past experience in the development and use of murine models of metastasis by our group. Each study was designed to minimize unnecessary animal use, optimize statistical power, and account for known variance in each model system. Within each experiment mice of the same strain, sex, and age were used for all conditions. At the initiation of each experiment mice were randomly assigned to cages and all mice in a given cage received equivalent treatment (e.g. doxycycline). Researchers were not blinded to the group assignments of mice as no subjective measurements were used.

### Human Subjects

Osteosarcoma primary and lung metastatic tumors were obtained from the Laboratory of Experimental Oncology, Rizzoli Institute, Bologna, Italy with approval from Rizzoli Institute Ethics Committee. A waiver was granted for informed consent for patients deceased at the time of data collection according to the Data Privacy Regulation. Estimated tumor cellularity for samples ranged from 50-90%.

Endometrial primary and metastatic tumors were obtained with protocol approval from University Hospitals Institutional Review Board (UHCMC IRB). UHCMC IRB has determined that with respect to the rights and welfare of the individuals, the appropriateness of the methods used to obtain informed consent, and the risks and potential medical benefits of the investigation, that the protocol is acceptable under Federal Human Subject Protection regulations promulgated under 45 CFR 46 and 21 CFR 50 and 56. Informed consent was obtained from all patients. The endometrial samples included a stage IIIC2 Uterine Serous Carcinoma and aortic lymph node metastasis. Estimated tumor cellularity was 40% and 90% for the primary and metastatic sample, respectively.

### *Ex Vivo* Lung Metastasis Assay

#### Procedure for RNA Isolation

GFP-positive tumor cells (5 x 10^5^) were delivered by tail vein injection to 8-10 week old female SCID/Beige (Charles River). Within 15 minutes of tumor cell injection, mice were euthanized with CO_2_ inhalation, and lungs were insufflated with a mixed agarose/media solution. Lung sections for *ex vivo* culture were generated as described^29^ and incubated at 37°C in humidified conditions of 5% CO_2_. Culture media was changed and lung sections were flipped every 2 days. Tumor cell RNA was harvested at 24hr and 14 day time points from one mouse for each condition. Lung sections were chopped into fine pieces and incubated in 3ml HBSS with 1mg/ml collagenase at 37°C for 30 minutes. EDTA was added to a final concentration of 10mM and the solution was placed on ice to stop digestion. Digested material was homogenized by passing through 18 ga needle 3-5x using 10 ml syringe. Homogenate was passed through a 70 micron cell strainer (Corning Life Sciences) and centrifuged at 500x*g* for 5 minutes at 4°C. Supernatant was aspirated and cells were re-suspended in 5ml ACK lysing buffer for 3 minutes at RT to lyse RBCs. Lysis was stopped by adding 10ml HBSS and cells were centrifuged at 500x*g* for 5 minutes at 4°C. Supernatant was aspirated and cells were re-suspended in 2-3ml 0.5mM EDTA PBS and placed on ice. Immediately prior to sorting, cells were passed through a 40 micron cell strainer (Corning Life Sciences).

Cells were sorted by FACS to isolate GFP+ tumor cells and immediately centrifuged at 500x*g* for 5 minutes at 4°C. Supernatant was aspirated and cells were lysed in 1ml TRIzol reagent (Life Technologies), RNA was extracted with 200ul chloroform. Organic phase was isolated and 700ul of EtOH added. RNA was purified from this solution using the RNeasy Micro Kit (Qiagen).

#### Procedure for F3 Knockdown Studies

F3 knockdown cells were pretreated with 5ug/ml doxycycline for 40hrs in standard culture. F3 shRNA induced cells were sorted by FACS to isolate DsRed+/GFP+ fraction. Uninduced control cells were sorted to isolate GFP+ fraction. 2x10^5^ cells were injected into the tail vein of each mouse. For F3 knockdown condition 5ug/ml doxycycline was added to agarose/media solution used to insufflate lungs as well as culture media. The medium was changed and fresh doxycycline was added every 2 days. A total of 8 lung sections were imaged for each condition (4 sections per mouse, 2 mice per condition).

#### Procedure for JQ1 Studies

5x10^5^ tumor cells were injected into the tail vein of each mouse. For JQ1 treated cultures, medium was supplemented to a final concentration of 250nM JQ1 by adding 10mM DMSO stock solution. Vehicle treated culture media was supplemented with DMSO volumes matching JQ1 treatment. Media was changed and fresh JQ1 or DMSO was added every 2 days. A total of 8 lung sections were imaged for each condition (4 sections per mouse, 2 mice per condition).

#### Assessment of Metastatic Burden

Lung sections were imaged by inverted fluorescent microscopy (Leica DM IRB) at a magnification of 2.5x. 2-3 images per lung section were taken to capture the entire surface of each section. Image analysis was performed using ImageJ software to quantify total GFP+ area per lung section. The metastatic burden was calculated by normalizing total GFP+ area to GFP+ area for each section on day 0. Values reported represent mean normalized tumor burden for all sections for each condition (8 sections per condition).

### *In Vitro* RNA isolation

To match conditions used to isolate cells from *ex vivo* lung sections, cells growing *in vitro* were trypsinized and exposed to the same mechanical/enzymatic digestion conditions and sorted by FACS as described above. 5x10^5^ GFP+ cells were collected and RNA was isolated as described above.

### ChIP-seq

ChIPs were performed from 5-10x10^6^ cross-linked cells and sequencing libraries were prepared as previously described^48^. The following antibodies were used for ChIP: rabbit anti-H3K4me1 (Abcam #8895), rabbit anti-H3K27ac (Abcam #4729), rabbit anti-c-Fos (Santa Cruz #sc-52), rabbit anti-FOSL1/Fra-1 (Santa Cruz #sc605). ChIP-seq libraries were sequenced on the HiSeq 2000 or 2500 platform at the Case Western Reserve University Genomics Core Facility.

The FASTX-Toolkit (http://hannonlab.cshl.edu/fastx_toolkit/) was used to remove adapter sequences and trim read ends using a quality score cutoff of 20. ChIPseq data were aligned to the hg19 genome assembly (retrieved from http://cufflinks.cbcb.umd.edu/igenomes.html), using Bowtie 2 v2.0.6^49^, allowing reads with ≤ 2 mismatches and discarding reads with > 1 reportable alignment (“- m 1” parameter). PCR duplicates were removed using SAMtools^50^. Peaks were detected with MACS v1.4^51^, using an aligned input DNA sample as control with a threshold for significant enrichment of P<1E-9. Wiggle tracks stepped at 25 bp were generated by MACS, normalized for the number of aligned reads and visualized on the UCSC Genome Browser.

### Met-VEL Analysis

H3K4me1 ChIP-seq peaks were filtered to remove all peaks overlapping ENCODE blacklisted regions for functional genomics analysis (https://sites.google.com/site/anshulkundaje/projects/blacklists) as well as peaks +/-1kb from transcription start sites (TSSs) of all annotated RefSeq genes to exclude promoters. Resulting peak lists of parental and metastatic cell line pairs were merged and RPKM values within merged peaks were calculated. Gained and lost Met-VELs were called as peaks with 3-fold increased or decreased RPKM values in metastatic cell lines relative to parental cell lines, respectively. To determine the fraction of differentially active enhancers in different cell types (Extended Figure 2), H3K4me1 ChIP-seq peaks for each pair of samples were filtered for ENCODE blacklisted regions and promoters, concatenated, and merged. Peak RPKMs were calculated for each sample in a pair and floored to 0.3. Differentially active enhancers were defined as those showing a 3-fold change in H3K4me1 signal in one sample relative to the other. The fractions of differentially active enhancers for the osteosarcoma tumors and cell lines panels were based on averages for each group.

### Met-VEL Clustering Analysis

Global Met VEL distribution was assessed by calculating Met-VEL counts in 200kb sliding windows across all chromosomes. Met-VEL islands were defined as regions bordered by 200kb windows with Met-VEL counts of 0. 200kb windows with maximum Met-VEL counts in each Met-VEL island were identified. To test for non-random Met-VEL distribution, the same analysis was performed on 1000 Met-VEL-size-matched H3K4me1 peak lists randomly sampled from all H3K4me1 peaks in the cell line being analyzed to account for global enhancer distribution biases. Metastatic cell line H3K4me1 peaks were sampled to assess gained Met-VEL clustering. Parental cell line H3K4me1 peaks were sampled to assess lost Met-VEL clustering. The sampled lists were used to define expected distributions of random VEL acquisition in each cell line. Expected distributions were compared to observed distributions to test the null hypothesis of random Met-VEL acquisition. A p-value threshold of 0.05 was used to reject the null hypothesis in support of non-random acquisition of Met-VELs. 200kb windows with Met-VEL counts exceeding these thresholds were called as Met-VEL clusters.

### Super-Enhancers (SEs)

Metastatic and parental cell line-specific SEs were identified using the ROSE and dynamicEnhancer software (retrieved from https://github.com/BradnerLab/pipeline). In each parental/metastatic cell line pair, H3K4me1 peaks identified by MACS were filtered to remove peaks within 2kb of RefSeq TSSs. Peaks separated by less than 12.5kb were stitched together. All stitched peaks were then ranked by the density of H3K4me1 minus input. Peaks higher than the inflection point on the density curve were designated SEs. To call unique SEs, SEs from parental and metastatic cell lines were merged and H3K4me1 signals for all merged SEs in each cell line were calculated. 2-fold change in H3K4me1 signal was used to define SEs unique to parental or metastatic cell lines.

### RNA-seq

Gene expression profiles of cell lines grown *in vitro* were compared to expression profiles of the same cell lines at various time points during metastatic colonization using the *ex vivo* pulmonary metastasis assay. RNA quality was assessed by 2200 TapeStation Instrument (Agilent). PolyA+ RNA was prepared for sequencing using the Illumina TruSeq RNA Sample Preparation Kit according to the manufacturer’s protocol. RNA-seq libraries were sequenced on the Illumina HiSeq 2000 or 2500 platform at the Case Western Reserve University Next Generation Sequencing Core Facility.

For gene expression analysis, reads were aligned to the hg19 genome build (retrieved from http://cufflinks.cbcb.umd.edu/igenomes.html), using Tophat v1.3.2^52^. FPKM values for known genes were calculated using Cufflinks v1.3.0^53^ provided with the GTF file via the –G (known genes only) option. Bias correction was used (-b option with hg19 FASTA as the input) to improve the accuracy of transcript abundance estimates. FPKM values were tabled by converting background values (<0.3) to 0 and adding 0.3 to all values^54^. FPKMs were quantile normalized across all samples.

### Prediction of Gene Targets of Enhancers Using PreSTIGE

Enhancer-gene assignments were made as described in^55^. Briefly, predictions were made using comparative analysis across a panel of 13 tissues. For an interaction to be predicted the normalized H3K4me1-enhancer signal intensity had to be above background and highly specific to the cell line of interest compared to the remaining 12 cell lines. Additionally, the gene must be within 100-kb of the enhancer and must show relatively cell type-specific transcript levels.

### Gene Ontology Analysis

#### Gained Met-VEL Gene Lists

Met-VEL gene lists were imported into gProfiler^56^ to generate enrichment scores for all GO, KEGG and REACTOME gene sets according to recommended settings for gProfiler http://baderlab.org/Software/EnrichmentMap/GProfilerTutorial. Cytoscape(v3.2.1) and the Enrichment Map^57^ plug-in was used to generate networks for gene sets enriched with an FDR cutoff of < 0.05.

#### Lost Met-VEL Gene Lists

For gene ontology (GO) analysis, the genes associated with Met-VELs were analyzed using DAVID (http://david.abcc.ncifcrf.gov/home.jsp). A p-value of 10^−3^ was used as the threshold for significant enrichment of an ontologic category. Categories significantly enriched for gained or lost Met VEL genes in 2 or more pairs are reported, limiting overlapping lists to the three top scoring categories in each cell line (i.e. the categories with the lowest p-values).

### DHS-seq

6.4-56x10^6^ cells from each cell line were subjected to DNase hypersensitivity (DHS) sequencing as previously described^58^ with the exception of a 5’ phosphate added to linker 1B to increase ligation efficiency. After DNase concentrations were optimized for each line a total of approximately 1x10^6^ cells from optimally digested conditions were processed for sequencing. Libraries were sequenced on the HiSeq 2500 platform at the Case Western Reserve University Genomics Core Facility.

Adapters and low quality bases (Phred score <20) were trimmed from the end of reads using the FASTX-Toolkit. Reads were aligned to the hg19 genome assembly using Bowtie 2 v2.0.6, allowing no more than 2 mismatches and discarding reads with > 1 reportable alignment (“-m 1” parameter). SAMtools was used to remove PCR duplicates. Peaks were identified using MACS v1.4. Tracks were visualized as described for ChIP-seq data.

### Chromosome Conformation Capture Sequencing (4C-seq)

4C-seq sample preparation was performed as previously described^59^. NlaIII served as a primary restriction enzyme, DpnII as a secondary 4 bp-cutter. Primer sequences are provided in below. Amplified sample libraries were pooled and spiked with 40% PhiX viral genome sequencing library to increase sample diversity. Multiplexed sequencing was performed on the MiSeq platform. Demultiplexing was performed by an in-house algorithm and all reads were hard trimmed to 36bp. Clipping of the primer sequences and data processing was performed using 4Cseqpipe Version 0.7 (retrieved from http://compgenomics.weizmann.ac.il/tanay/?page id=367). The viewpoint reads were aligned to a fragmented genome, as determined by the restriction site positions of the chosen primary and secondary restriction enzymes. A running linearly weighted mean, calculated in sliding windows of size 2-50KB, was used for signal smoothing of each genomic bin (size 16bp). Contact enrichment sites along the chromosomal axis were visually inspected.

**Table 2-1:**
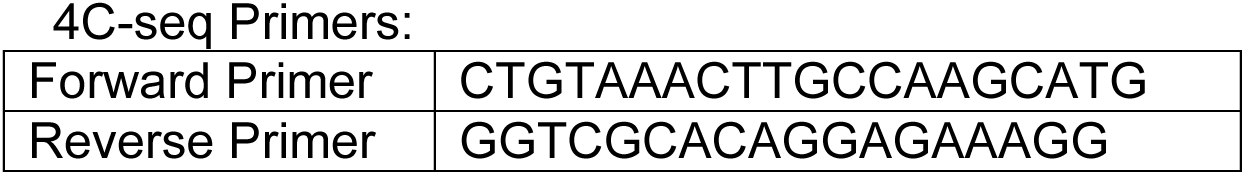
4C-seq primers.

### Motif Analysis

To identify transcription factor (TF) motifs enriched in Met-VEL peaks, enhancers were centered on DNase hypersensitivity sites and the SeqPos module of the Cistrome tool was used to scan a 1kb window for enriched curated motifs^60^. Significantly enriched motifs in each cell line were then filtered using RNA-seq data and only expressed TFs were used for downstream analysis. Expressed TFs with enriched motifs in 3 out of 3 metastatic/parental cell line pairs (MG63.3/MG63; MNNG/HOS; 143B/HOS) are presented in the results.

### *In Vivo* RNAi High-Throughput Functional Assay

#### Vector Construction

The Tet-ON lentiviral construct was made by modifying the previously published optimized shRNAmir, “miR-E”, pRRL backbone^34^. Briefly, this construct contains an optimized 3^rd^ generation Tet-responsive element (T3G) and rtTA3 to potentiate a positive feedback loop, enhancing expression of the construct upon induction and reducing construct leakiness. The version of the construct that we modified contained a constitutive Venus reporter and an induced DsRed reporter of expression (LT3REVIR). The construct was modified using standard cloning techniques to replace the Venus reporter with a puromycin resistance element (renamed LT3REPIR) so that cells already constitutively expressing GFP could be selected for transduction.

#### shRNA Library Generation

shRNAs targeting 33 genes were selected from the transOMIC technologies shERWOOD-UltramiR shRNA library (3 to 4 shRNAs per gene). Cloning of shRNA into the backbone construct was performed on contract by transOMIC technologies. The following shRNA sequences were included in the library:

Scores indicate shERWOOD metric of predicted potency of each shRNA as assigned by previously algorithm^61^ described. NGS of #N/A indicates that the shRNA failed to clone into the lentiviral backbone.

**Table 2-2:**
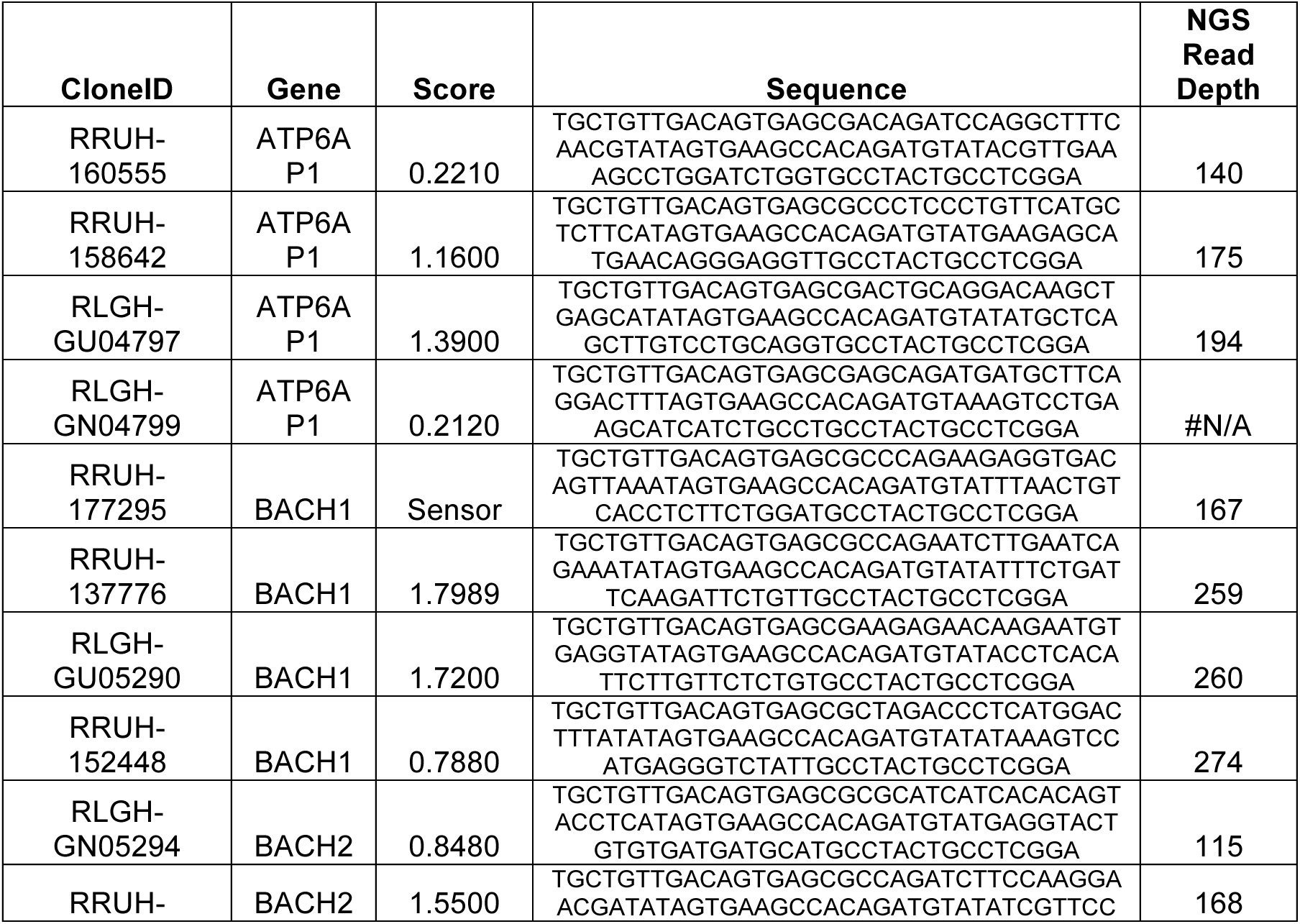
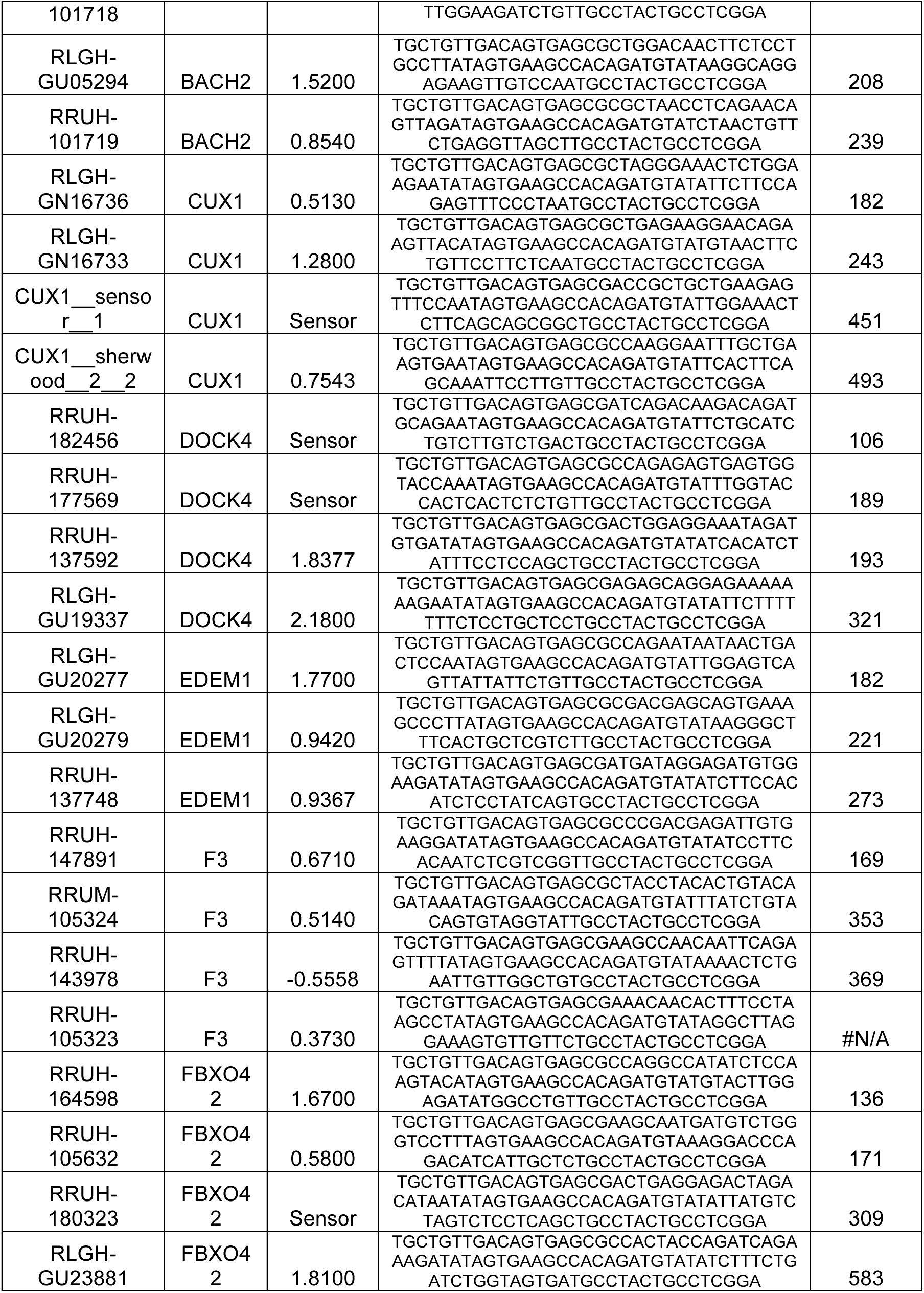
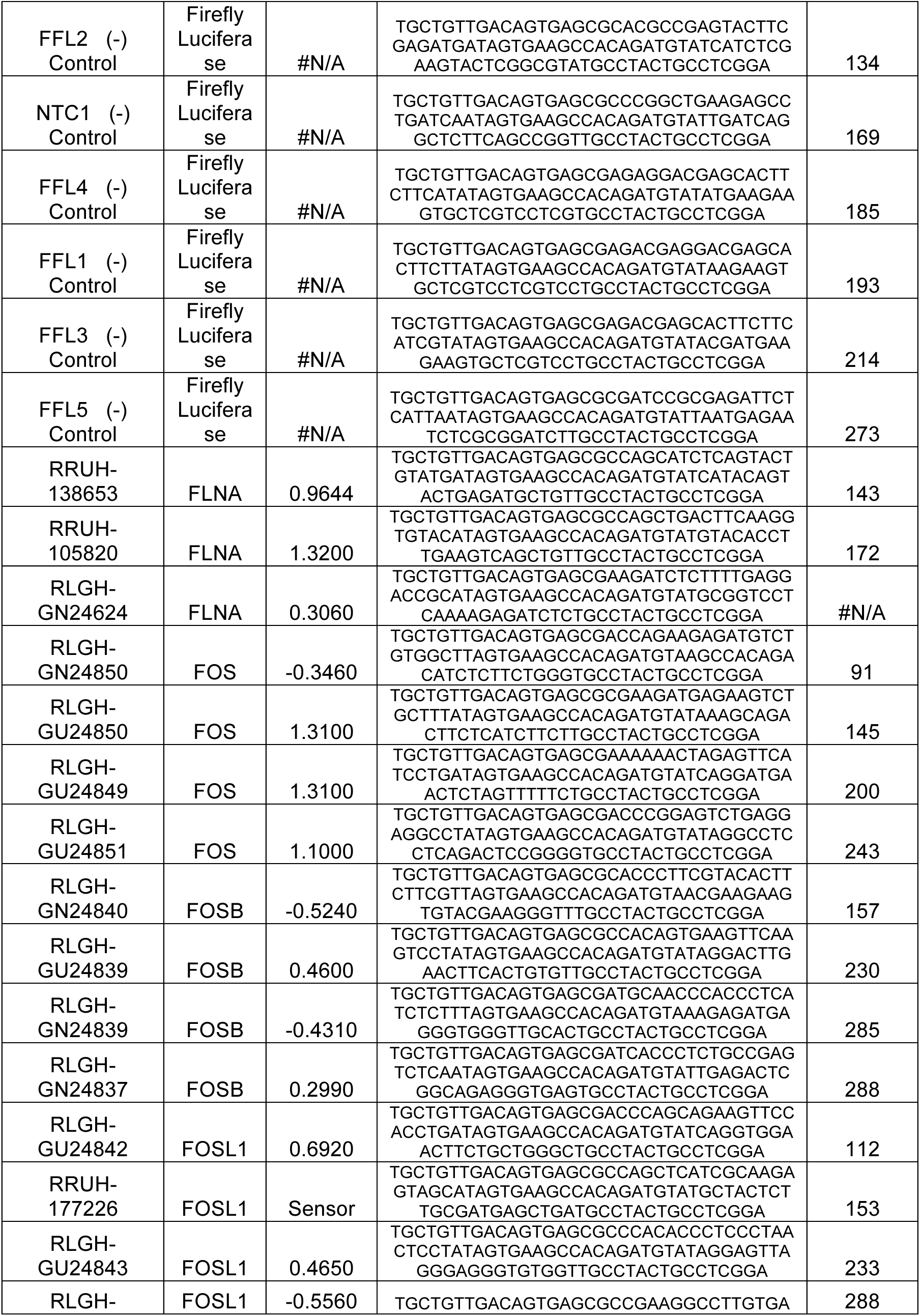
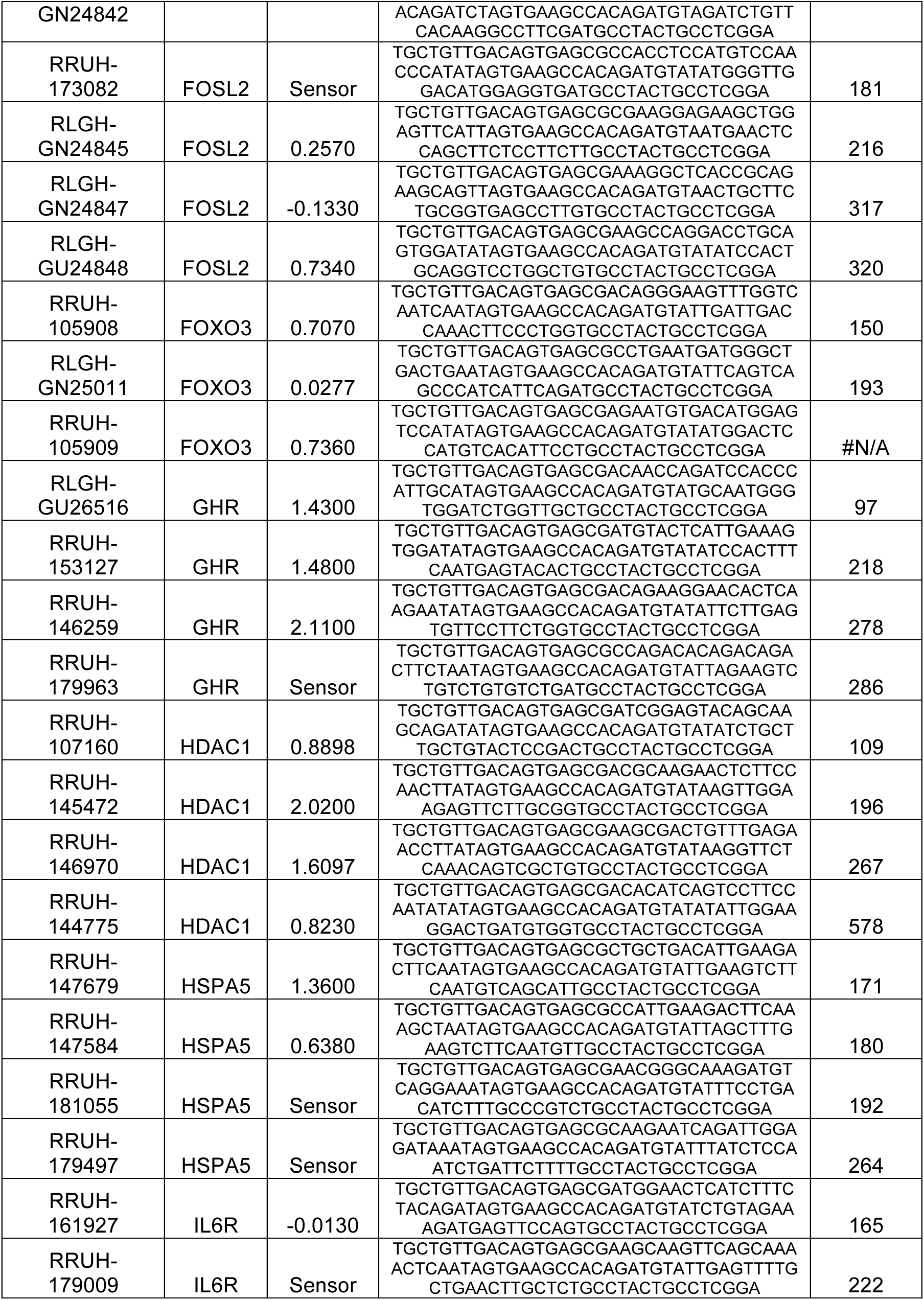
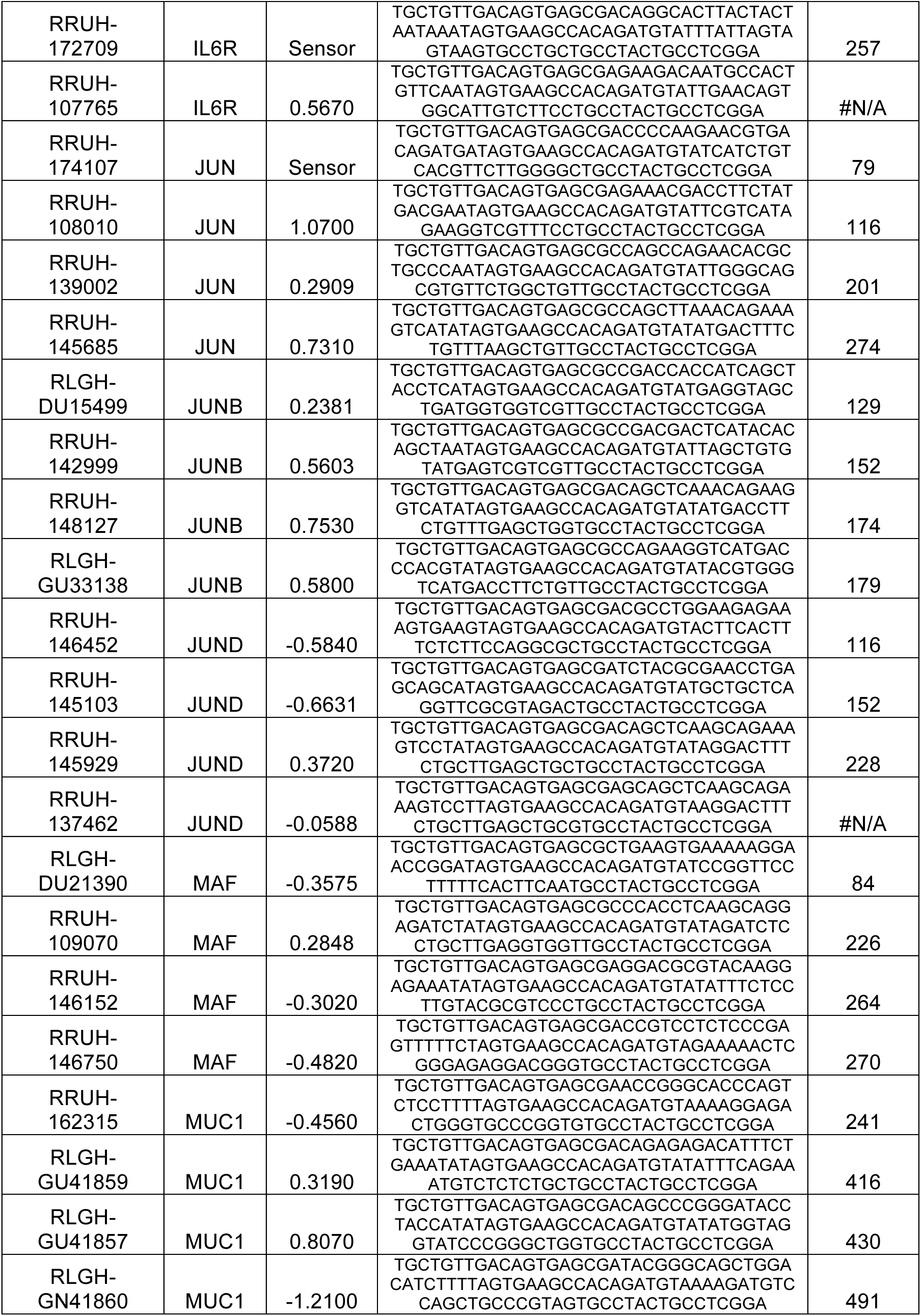
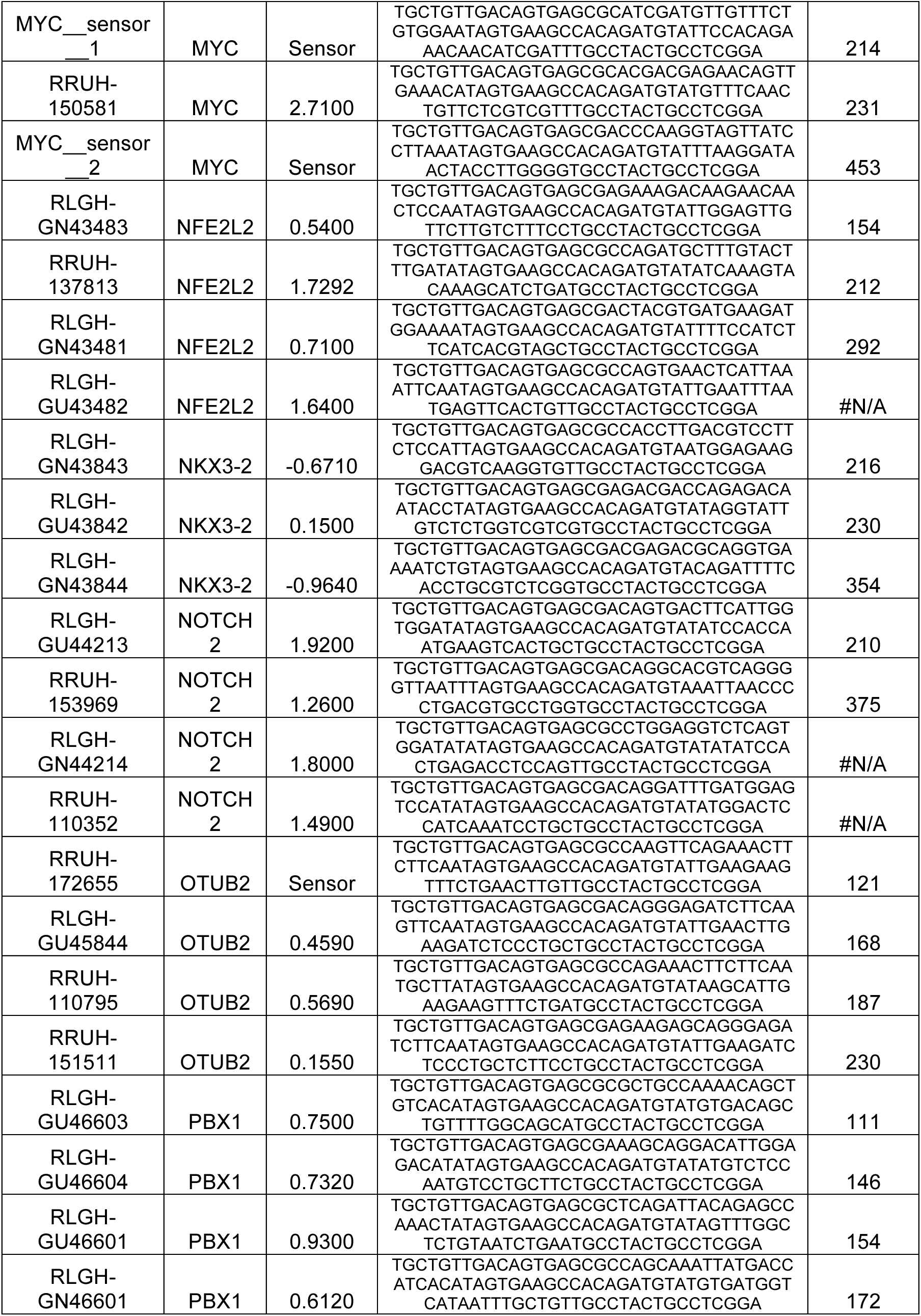
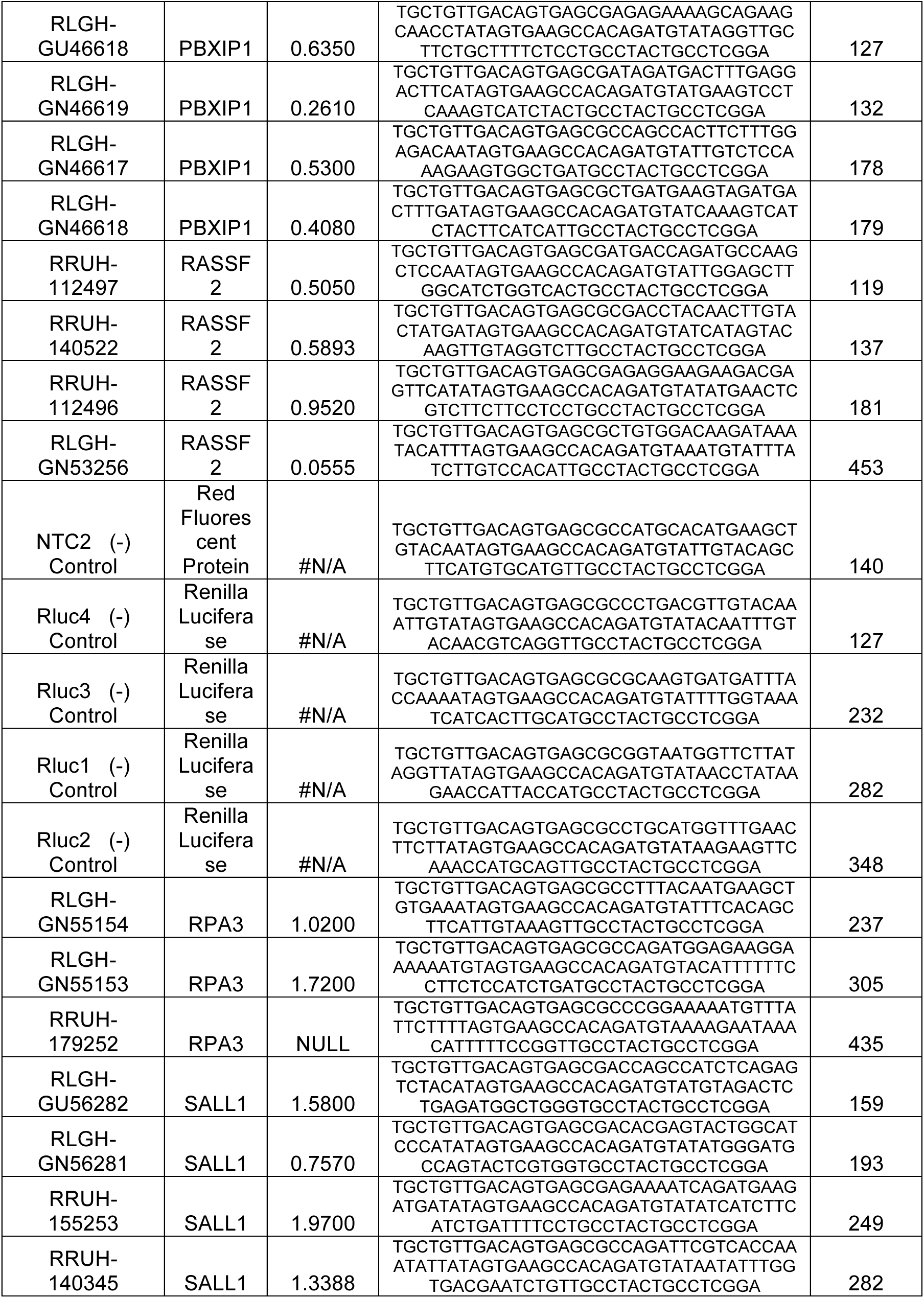
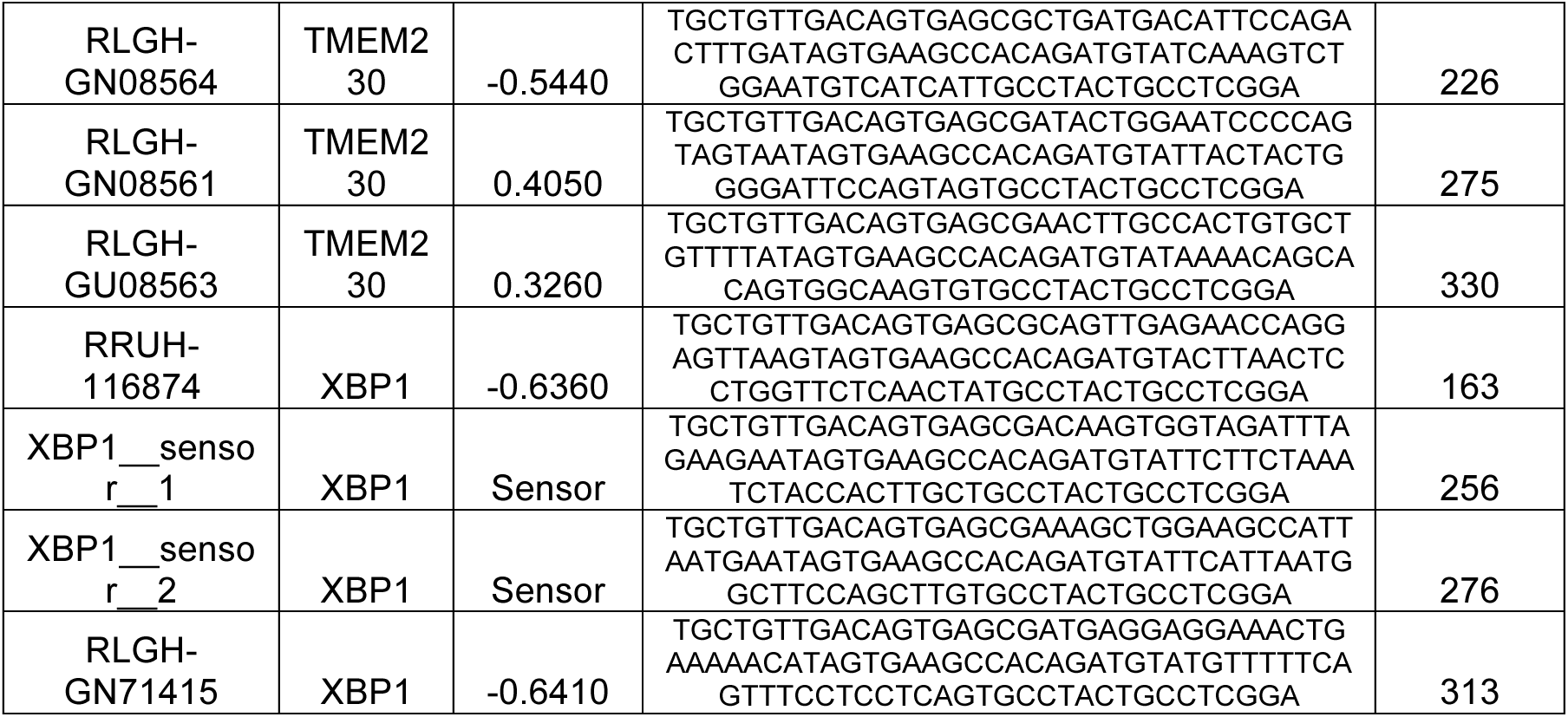
Hairpins used in high throughput *in vivo* RNAi screen.

#### Lentiviral Production

VSV-G pseudotyped lentivirus was generated with standard laboratory techniques. Briefly, shRNA-LT3REPIR plasmids were co-transfected with packaging vectors psPAX2 and pCI-VSVG (Addgene) into 293FT cells using Cal-Phos Mammalian Transfection Kit (Clontech). Individual supernatants containing virus were harvested at 48 and 72 h post-transfection and filtered with 0.45μm PVDF membrane (Millipore).

#### Lentiviral Transduction and Selection

Transduction of MG63.3 was performed via 24hr exposure to lentivirus in the presence of 8ug/ml polybrene using conditions to achieve >1000x coverage of each shRNA in the library. Infection rate estimated to be 0.135% and predicted to achieve predominantly one lentiviral integration per cell. Transduced and nontransduced cells were then treated with 2ug/ml puromycin. Transduced cells were selected until all cells in non-transduced plate were dead (2-4days) to obtain a pure population of transduced cells (MG63.3i).

#### High-Throughput In Vivo Functional Assay

8-10 week old female SCID/Beige (Charles River) mice were used for the *in vivo* study arm. Mice were fed Dox Diet pellets containing 200mg/kg doxycycline (Bio-Serve) for 5 days prior to injection of cells. MG63.3i cells were pre-treated with 5ug/ml doxycycline for 12hrs in standard culture before being delivered to mouse lungs by tail vein injection. This primes the cells, but knockdown is not achieved even at the transcript level until 24-72 h after doxycycline addition. 1.5x10^6^ cells were injected into each mouse (n=15). Mice were maintained on Dox Diet throughout the 21-day course of the experiment. At the conclusion of the experiment mice were euthanized by CO_2_ inhalation and lungs were surgically extracted and homogenized using the Tumor Dissociation Kit, human (Miltenyi) according to the manufacturer’s protocol. Mouse lung cells were depleted using the Mouse Cell Depletion Kit (Miltenyi) according to the manufacturer’s protocol. Lungs from 5 mice were pooled for each replicate to achieve 1000x engraftment coverage of each shRNA in the library. GFP+/DsRed+ cells were then isolated by FACS.

Three replicates of 1.5x10^6^ MG63.3i cells growing *in vitro* were induced with 5ug/ml doxycycline and maintained on doxycycline over 21 days in culture. GFP+/DsRed+ cells were isolated by FACS. Sorted cell counts of *in vitro* replicates were matched to numbers of cells isolated from the *in vivo* arm (3- 5x105).

DNA was isolated from three replicates of uninduced MG63.3i cells as well as *in vivo* and *in vitro* arms of the experiment.

#### shRNA Amplification and Sequencing

Genomic DNA was isolated and sequenced as described^62^ with slight modification. Genomic DNA was isolated by two rounds of phenol extraction using PhaseLock tubes (5prime) followed by isopropanol precipitation. Deep sequencing libraries were generated by PCR amplification of shRNA guide strands using barcoded primers that tag the product with standard Illumina adapters. For each sample, DNA from at least 3 × 10^5^ cells was used as template in multiple parallel 50-μl PCR reactions, each containing 1 μg template, 1× AmpliTaq Gold buffer, 0.2 mM of each dNTP, 0.3 μM of each primer and 2.5 U AmpliTaq Gold (Applied Biosystems), which were run using the following cycling parameters: 95 °C for 10 min; 35 cycles of 95 °C for 30 s, 52 °C for 45 s and 72 °C for 60 s; 72 °C for 7 min. PCR products (340 nt) were combined for each sample, precipitated and purified on a 2% agarose gel (QIAquick gel extraction kit, Qiagen). Libraries were sequenced on the HiSeq 2500 platform at the Case Western Reserve University Genomics Core Facility. Libraries were sequenced using a primer that reads in reverse into the guide strand (miR30EcoRISeq, TAGCCCCTTGAATTCCGAGGCAGTAGGCA). To provide a sufficient baseline for detecting shRNA depletion in experimental samples, we aimed to acquire >1,000 reads per shRNA in all samples or 1.37x10^5^ reads per sample. In practice, we achieved >2x10^6^ reads for all samples. Sequence processing was performed using two custom workflows using usegalaxy.org^63^. Workflow can be accessed by the following links: https://usegalaxy.org/u/tyleremiller/w/shrna-pipeline1 and https://usegalaxy.org/u/tyleremiller/w/shrnastep2. For each shRNA and condition, the number of matching reads was normalized to the total read number per lane. This measure of normalized coverage was used for all downstream analyses.

### Inducible Knockdown of *Tissue Factor (F3)*

shRNAs targeting *tissue factor (F3)* were selected from the transOMIC technologies shERWOOD-UltramiR shRNA library and cloned into the LT3REPIR as described in the preceding section. Cloning of shRNA into the backbone construct was performed on contract by transOMIC technolgies. The following shRNA sequences were tested:

**Table 2-3:**
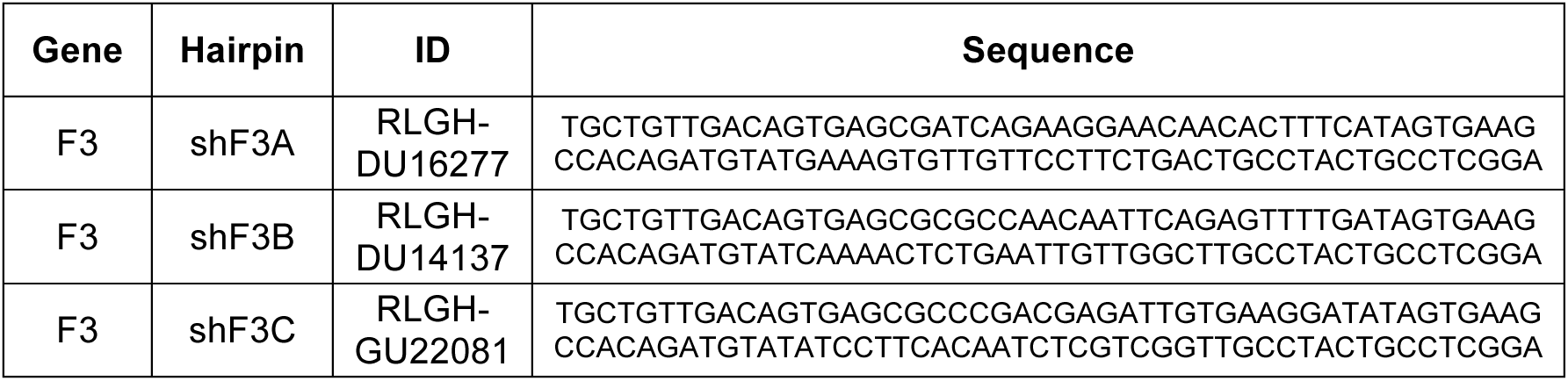
Hairpins used in *F3* knockdown experiments.

#### Lentiviral Production

VSV-G pseudotyped lentivirus was generated with standard laboratory techniques. Briefly, shRNA-LT3REPIR plasmids were cotransfected with packaging vectors psPAX2 and pCI-VSVG (Addgene) into 293FT cells using Cal-Phos Mammalian Transfection Kit (Clontech). Individual supernatants containing virus were harvested at 48 and 72 h post-transfection and filtered with 0.45 μm PVDF membrane (Millipore).

#### Lentiviral Transduction and Selection

Transduction was performed via 24hr exposure to lentivirus in the presence of 8ug/ml polybrene. Transduced and non-transduced cells were then treated with 2ug/ml puromycin. Transduced cells were selected until all cells in non-transduced plate were dead (2-4days).

#### Assessment of Knockdown

Optimal shRNA induction was assessed and found to occur with 5ug/ml doxycycline treatment. MG63.3 cells transduced with shF3A, shF3B, and shF3C were treated with doxycycline for 40hrs, typsinized and sorted to isolate DsRed+/GFP+ fraction. Uninduced cells were sorted for GFP+ fraction. RNA was extracted from 1x10^6^ cells and purified using the RNAeasy Micro kit (Qiagen) according to the manufacturer’s protocol. RNA quality was assessed by 2200 TapeStation Instrument (Agilent). cDNA was synthesized using the High Capacity RNA-to-cDNA kit (ABI) according to the manufacturer’s protocol. Knockdown efficiency was determined by RT-qPCR for *F3* using optimized TaqMan Gene Expression Assay primers and TaqMan Gene Expression Master Mix (Life Technologies).

**Table 2-4:** Probes used for *F3* RT-qPCR. RT-qPCR was performed to quantify percent knockdown with induction of each hairpin relative to uninduced controls. Hairpins shF3A and shF3B showed the highest degree of *F3* knockdown and were chosen for use the remainder of these studies.

| Gene Symbol | TaqMan Gene Expression Assay Primer |
| --- | --- |
| *F3* | Hs01076029_m1 |
| *GAPDH* | Hs02758991_g1 |

### *In Vivo* Experimental Metastasis Model

8-10 week old female SCID/Beige (Charles River) mice were used for all experimental metastasis studies. For all *in vivo* experimental *F3* knockdown studies, mice in the *F3* knockdown group were fed Dox Diet pellets containing 2gm/kg doxycycline (Bio-Serve) for 5 days prior to injection of cells. *F3* knockdown cells were pre-treated with 5ug/ml doxycycline for 24hrs in standard culture. *F3* shRNA induced cells were sorted by FACS to isolate DsRed+/GFP+fraction. Control cells were sorted to isolate GFP+ fraction.

#### End Point Assessment of Lung Metastasis

5x10^5^ MG63.3 DsRed+/GFP+ or MG63.3 GFP+ cells were injected into the tail vein of each mouse (n=10 mice per condition). Mice in the F3 knockdown group were maintained on Dox Diet throughout the experiment. On day 7 or day 14 following injection 5 mice from each group were euthanized by CO_2_ inhalation. Lungs were insufflated with PBS and imaged by inverted fluorescent microscopy (Leica DM IRB) at a magnification of 2.5x. 5 images per lung were taken to assess metastatic burden in each mouse.

Image analysis was performed using ImageJ software to quantify total GFP+ area per image. The metastatic burden was calculated as the sum of the total GFP+ area in the 5 images from each mouse.

#### Survival Analysis

5x10^4^ cells were injected into the tail vein of each mouse (n=5 mice per condition). Mice in the F3 knockdown group were maintained on Dox Diet throughout the experiment. All mice that died underwent complete necropsy examination and confirmation of metastasis.

### Orthotopic Spontaneous Lung Metastasis Model

8-10 week old female NSG mice (Jackson) were used for spontaneous metastasis studies. Mice in the F3 knockdown group were given water supplemented with 2mg/ml doxycycline hyclate (Sigma) and 2% sucrose for 5 days prior to injection of cells. F3 knockdown cells were pre-treated with 5ug/ml doxycycline for 24hrs in standard culture. F3 shRNA induced cells were sorted by FACS to isolate DsRed+/GFP+ fraction. Control cells were sorted to isolate GFP+ fraction. 3x10^5^ cells were injected orthotopically into the paraosseous region adjacent to the left proximal tibia. For the F3 knockdown group water was changed and fresh doxycycline was added twice weekly. Injection sites were monitored twice weekly for tumor formation.

Tumors became measureable on day 21 for all groups at which time tumors were measured in two dimensions twice weekly. Tumor volume was calculated as follows: volume (mm^3^) = 3.14 x [long dimension (mm)] x [short dimension (mm)]^2^. Experiment was terminated following 21 days of tumor measurements (42 days after injections) and mice were euthanized by CO_2_ inhalation. Lungs were insufflated with PBS and imaged by fluorescent microscopy at 2.5x using Leica DM 5500B light microscope with a Leica DFC 500 camera. 5 images per lung were taken to assess metastatic burden in each mouse.

Image analysis was performed using ImageJ software. Micrometastases were defined as GFP+ lesions with diameter >25 pixels in images captured at 2.5x magnification. Number of micrometastases per image were manually counted. Total micrometastases for each lung were calculated as the sum of the total number of micrometastases in 5 images from each mouse.

### *In Vitro* Assay for FXa Formation

Cells growing *in vitro* were pre-treated with 25μg/mL IgG control, Mab-10H10, or Mab-5G9 20 minutes prior to assay. Cells were washed in serum-free DMEM, and Xa generation over time was measured 30 minutes after the addition of 1nM FVIIa and 50nM FX using the chromogenic substrate Spectrozyme FXa.

### Assessment of F3 Inhibiting Antibodies on Metastatic Progression

5x10^5^ GFP+ MG63.3 cells were mixed with 500μg IgG control, Mab-10H10, or Mab-5G9 and injected into the tail vein of 10-12 week old female SCID-beige mice (N>5 mice per group). Mice were sacrificed 14 days after injection and metastatic burden was assessed by whole lung fluorescent imaging (5 images per mouse). Metastatic burden was quantified as total GFP+ area per mouse.

### F3 Lung Metastasis Staining

To assess the F3 expression in metastatic tumor cells in the lung at progressive time points, mice were injected with 1 x 10^6^ cells (via tail vein) and were euthanized via CO2 inhalation at 24 hrs and 15 days post-injection. Lungs were harvested, formalin-fixed and paraffin embedded. For *ex vivo* lung metastasis staining, the protocol described above was followed and lung sections were fixed at 24hrs and 5 days post-injection. Tissue sections of lungs were cut at a thickness of 5 microns. Prior to immunostaining, paraffin sections were dewaxed with xylenes, and rehydrated with an ethanol series. For antigen retrieval, tissue sections were immersed 95°C Target Retreival Solution (DAKO) for 25 minutes. Tissue sections were permeabilized with 0.01% Triton-X in PBS for 10 minutes. Slides were rinsed with PBS and blocked with 4% BSA in PBS for 10 minutes. The following primary antibodies were used: F3 - Rabbit monoclonal IgG F3 antibody (ab151748, Abcam); GFP – Goat polyclonal IgG GFP antibody conjugated to FITC (ab6662, Abcam). Primary antibodies were diluted in 4% BSA 1:100 and slides were incubated in antibody solution at 4°C overnight. Slides were rinsed and incubated with goat polyclonal IgG anti-rabbit IgG (H+L) conjugated to Alexa 594 (A-11037, Life Technologies) diluted 1:200 in 4% BSA for 1hr in dark humidified slide chamber. Nuclei were visualized with DAPI (Sigma, 1ug/ml). Tissue sections were mounted on slides using anti-fade mounting medium (Vectashield).

Stained sections were imaged by fluorescent microscopy at 20 or 40x using Leica DM 5500B light microscope with a Leica DFC 500 camera. Image analysis was performed using ImageJ software. F3 expression was computed within GFP+ metastatic tumor cell area.

### Tissue Array Staining and Scoring

A tissue microarray that we previously developed containing 20 osteosarcoma patient lung metastases^64^ was assessed for F3 expression. 18 of these 20 samples had cores of sufficient quality on the stained slide to be scored. For staining, paraffin was removed by 5 minute incubation in xylene bath x2 and rehydrated using step-down concentrations of EtOH. Antigen retrieval was performed by incubation in 1:10 dilution of Target Retrieval Solution (DAKO) in steamer for 25 minutes at 95°C. Cells were permeabilized with 0.01% Triton-X in PBS for 10 minutes. F3 immunohistochemical staining was performed using rabbit monoclonal IgG F3 antibody (ab151748, Abcam) and the EnVision+ System-HRP (Dako) according to the manufacturer’s protocol. Cover slips were mounted on slides using anti-fade mounting medium.

The array was scored by the Director of Soft Tissue Pathology at the Cleveland Clinic and Learner Research Institute who was blinded to the sample type. Cores were scored based on the % of tumor cells in the core with positive staining for F3 (0=0%, 1+=1-25%, 2+=26-50%, 3+=>50%) and the intensity of F3 staining in positive areas (low intensity staining, high intensity staining). Individual cores were excluded from the analysis if no tumor was present, tumor was predominantly necrotic, or core was falling off the slide.

### Targeted Deletion of an *F3* Met-VEL by Genome Engineering

Two TALEN dimers were designed to target the flanks of the Met-VEL in the *F3* locus as indicated in Fig. 5a. TALEN dimers recognized the sequences 5′-GACCAACTCACTTGAGCTGtgtggtttttcttCAGTGCACAATTGTGAAAT-3’ and 5′-GAATCGACTGATCAAAGCacatgaactttttaaaaaaGAGTAATAAGTTTACTT-3′, where spacer elements are in lower case. TALEN constructs were assembled with adaptations of previously described protocols^65, 66^. MG63.3 cells were grown in 6-well plate format to 70% confluence and transfected with 2.5 μg plasmid for each TALEN monomer using Lipofectamine 2000® (ThermoFisher Scientific) as per the manufacturer’s instructions. Cells were incubated for 48 h at 30°C and genomic DNA was subsequently harvested using QuickExtract™ DNA Extraction Solution (Epicentre) as recommended by the supplier. Efficient deletion of the *F3* Met-VEL was confirmed by agarose gel electrophoresis of PCR products generated using primers 5′-GCAGTGCACAACCTGTACAAC-3’ and 5′-TTGGCCAGGGTCATTATGTT-3’ (Integrated DNA Technologies) and high fidelity AccuPrime *Taq* DNA Polymerase (ThermoFisher Scientific). Single cell clones were derived by limiting dilution and genotypes were confirmed as described above. Enhancer deletion of clonal cell population used for functional metastasis experiments was confirmed by Sanger sequencing using the primer sequences listed above.

## References

1 Valastyan, S. & Weinberg, R. A. Tumor metastasis: molecular insights and evolving paradigms. Cell 147, 275–292, doi:10.1016/j.cell.2011.09.024 (2011).

2 Chambers, A. F., Groom, A. C. & MacDonald, I. C. Dissemination and growth of cancer cells in metastatic sites. Nature reviews. Cancer 2, 563–572, doi:10.1038/nrc865 (2002).

3 Gundem, G. et al. The evolutionary history of lethal metastatic prostate cancer. Nature 520, 353–357, doi:10.1038/nature14347 (2015).

4 Hong, M. K., Macintyre, G., Wedge, D. C. & Van Loo, P. Tracking the origins and drivers of subclonal metastatic expansion in prostate cancer. Nature communications 6, 6605, doi: 10.1038/ncomms7605 (2015).

5 Bos, P. D. et al. Genes that mediate breast cancer metastasis to the brain. Nature 459, 1005–1009, doi:10.1038/nature08021 (2009).

6 Kang, Y. et al. A multigenic program mediating breast cancer metastasis to bone. Cancer cell 3, 537–549 (2003).

7 Minn, A. J. et al. Genes that mediate breast cancer metastasis to lung. Nature 436, 518–524, doi:10.1038/nature03799 (2005).

8 Factor, D. C. et al. Epigenomic comparison reveals activation of “seed” enhancers during transition from naive to primed pluripotency. Cell stem cell 14, 854–863, doi:10.1016/j.stem.2014.05.005 (2014).

9 Gifford, C. A. et al. Transcriptional and epigenetic dynamics during specification of human embryonic stem cells. Cell 153, 1149–1163, doi:10.1016/j.cell.2013.04.037 (2013).

10 Zhu, J. et al. Genome-wide chromatin state transitions associated with developmental and environmental cues. Cell 152, 642–654, doi:10.1016/j.cell.2012.12.033 (2013).

11 Heintzman, N. D. et al. Histone modifications at human enhancers reflect global cell-type-specific gene expression. Nature 459, 108–112, doi:10.1038/nature07829 (2009).

12 Akhtar-Zaidi, B. et al. Epigenomic enhancer profiling defines a signature of colon cancer. Science (New York, N.Y.) 336, 736–739, doi:10.1126/science.1217277 (2012).

13 Cohen, A. J. et al. Hotspots of aberrant enhancer activity punctuate the colorectal cancer epigenome. Nature communications 8, 14400, doi:10.1038/ncomms14400 (2017).

14 Hnisz, D. et al. Super-enhancers in the control of cell identity and disease. Cell 155, 934–947, doi:10.1016/j.cell.2013.09.053 (2013).

15 Loven, J. et al. Selective inhibition of tumor oncogenes by disruption of super-enhancers. Cell 153, 320–334, doi:10.1016/j.cell.2013.03.036 (2013).

16 Ramaswamy, S., Ross, K. N., Lander, E. S. & Golub, T. R. A molecular signature of metastasis in primary solid tumors. Nat. Genet. 33, 49–54, doi:10.1038/ng1060 (2003).

17 McDonald, O. G. et al. Epigenomic reprogramming during pancreatic cancer progression links anabolic glucose metabolism to distant metastasis. Nat. Genet. 49, 367–376, doi: 10.1038/ng.3753 (2017).

18 Kansara, M., Teng, M. W., Smyth, M. J. & Thomas, D. M. Translational biology of osteosarcoma. Nature reviews. Cancer 14, 722–735, doi: 10.1038/nrc3838 (2014).

19 Huang, Y. M., Hou, C. H., Hou, S. M. & Yang, R. S. The metastasectomy and timing of pulmonary metastases on the outcome of osteosarcoma patients. Clinical medicine. Oncology 3, 99–105 (2009).

20 Whyte, W. A. et al. Master transcription factors and mediator establish super-enhancers at key cell identity genes. Cell 153, 307–319, doi:10.1016/j.cell.2013.03.035 (2013).

21 Ren, L. et al. Characterization of the metastatic phenotype of a panel of established osteosarcoma cells. Oncotarget (2015).

22 Zentner, G. E., Tesar, P. J. & Scacheri, P. C. Epigenetic signatures distinguish multiple classes of enhancers with distinct cellular functions. Genome research 21, 1273–1283, doi:10.1101/gr.122382.111 (2011).

23 Rada-Iglesias, A. et al. A unique chromatin signature uncovers early developmental enhancers in humans. Nature 470, 279–283, doi:10.1038/nature09692 (2011).

24 Huang, H., Bhat, A., Woodnutt, G. & Lappe, R. Targeting the ANGPT-TIE2 pathway in malignancy. Nature reviews. Cancer 10, 575–585, doi: 10.1038/nrc2894 (2010).

25 Clayton, P. E., Banerjee, I., Murray, P. G. & Renehan, A. G. Growth hormone, the insulin-like growth factor axis, insulin and cancer risk. Nat. Rev. Endocrinol. 7, 11–24, doi:10.1038/nrendo.2010.171 (2011).

26 Pinski, J. et al. Inhibition of growth of human osteosarcomas by antagonists of growth hormone-releasing hormone. J. Natl. Cancer Inst. 87, 1787–1794 (1995).

27 Li, N. et al. Phosp hod ieste rase 10A: a novel target for selective inhibition of colon tumor cell growth and beta-catenin-dependent TCF transcriptional activity. Oncogene 34, 1499–1509, doi:10.1038/onc.2014.94 (2015).

28 van den Berg, Y. W., Osanto, S., Reitsma, P. H. & Versteeg, H. H. The relationship between tissue factor and cancer progression: insights from bench and bedside. Blood 119, 924–932, doi:10.1182/blood-2011-06-317685 (2012).

29 Mendoza, A. et al. Modeling metastasis biology and therapy in real time in the mouse lung. The Journal of clinical investigation 120, 2979–2988, doi:10.1172/jci40252 (2010).

30 Leaner, V. D. et al. Inhibition of AP-1 transcriptional activity blocks the migration, invasion, and experimental metastasis of murine osteosarcoma. The American journal of pathology 174, 265–275, doi:10.2353/ajpath.2009.071006 (2009).

31 Lamoureux, F. et al. Selective inhibition of BET bromodomain epigenetic signalling interferes with the bone-associated tumour vicious cycle. Nature communications 5, 3511, doi: 10.1038/ncomms4511 (2014).

32 Puissant, A. et al. Targeting MYCN in neuroblastoma by BET bromodomain inhibition. Cancer discovery 3, 308–323, doi:10.1158/2159-8290.cd-12-0418 (2013).

33 Bandopadhayay, P. et al. BET bromodomain inhibition of MYC-amplified medulloblastoma. Clinical cancer research: an official journal of the American Association for Cancer Research 20, 912–925, doi:10.1158/1078-0432.ccr-13-2281 (2014).

34 Fellmann, C. et al. An optimized microRNA backbone for effective single-copy RNAi. Cell reports 5, 1704–1713, doi:10.1016/j.celrep.2013.11.020 (2013).

35 Versteeg, H. H. et al. Inhibition of tissue factor signaling suppresses tumor growth. Blood 111, 190–199, doi:10.1182/blood-2007-07-101048 (2008).

36 You, J. S. & Jones, P. A. Cancer genetics and epigenetics: two sides of the same coin? Cancer cell 22, 9–20, doi:10.1016/j.ccr.2012.06.008 (2012).

37 Jones, S. et al. Comparative lesion sequencing provides insights into tumor evolution. Proc. Natl. Acad. Sci. U. S. A. 105, 4283–4288, doi:10.1073/pnas.0712345105 (2008).

38 Liu, W. et al. Copy number analysis indicates monoclonal origin of lethal metastatic prostate cancer. Nat. Med. 15, 559–565, doi:10.1038/nm.1944 (2009).

39 Campbell, P. J. et al. The patterns and dynamics of genomic instability in metastatic pancreatic cancer. Nature 467, 1109–1113, doi:10.1038/nature09460 (2010).

40 Yachida, S. et al. Distant metastasis occurs late during the genetic evolution of pancreatic cancer. Nature 467, 1114–1117, doi:10.1038/nature09515 (2010).

41 Navin, N. et al. Tumour evolution inferred by single-cell sequencing. Nature 472, 90–94, doi:10.1038/nature09807 (2011).

42 Moelans, C. B. et al. Genomic evolution from primary breast carcinoma to distant metastasis: Few copy number changes of breast cancer related genes. Cancer Lett. 344, 138–146, doi:10.1016/j.canlet.2013.10.025 (2014).

43 Kerbel, R. S., Frost, P., Liteplo, R., Carlow, D. A. & Elliott, B. E. Possible epigenetic mechanisms of tumor progression: induction of high-frequency heritable but phenotypically unstable changes in the tumorigenic and metastatic properties of tumor cell populations by 5-azacytidine treatment. J. Cell. Physiol. Suppl. 3, 87–97 (1984).

44 Rodenhiser, D. I. Epigenetic contributions to cancer metastasis. Clinical experimental metastasis 26, 5–18, doi:10.1007/s10585-008-9166-2 (2009).

45 Javaid, S. et al. Dynamic chromatin modification sustains epithelial-mesenchymal transition following inducible expression of Snail-1. Cell reports 5, 1679–1689, doi:10.1016/j.celrep.2013.11.034 (2013).

46 Latil, M. et al. Cell-Type-Specific Chromatin States Differentially Prime Squamous Cell Carcinoma Tumor-Initiating Cells for Epithelial to Mesenchymal Transition. Cell stem cell 20, 191–204.e195, doi:10.1016/j.stem.2016.10.018 (2017).

47 Khanna, C. et al. An orthotopic model of murine osteosarcoma with clonally related variants differing in pulmonary metastatic potential. Clinical & experimental metastasis 18, 261–271 (2000).

48 Schmidt, D. et al. ChIP-seq: using high-throughput sequencing to discover protein-DNA interactions. Methods (San Diego, Calif.) 48, 240–248, doi:10.1016/j.ymeth.2009.03.001 (2009).

49 Langmead, B., Trapnell, C., Pop, M. & Salzberg, S. L. Ultrafast and memory-efficient alignment of short DNA sequences to the human genome. Genome biology 10, R25, doi:10.1186/gb-2009-10-3-r25 (2009).

50 Li, H. et al. The Sequence Alignment/Map format and SAMtools.Bioinformatics (Oxford, England) 25, 2078–2079, doi:10.1093/bioinformatics/btp352 (2009).

51 Zhang, Y. et al. Model-based analysis of ChIP-Seq (MACS). Genome biology 9, R137, doi:10.1186/gb-2008-9-9-r137 (2008).

52 Trapnell, C., Pachter, L. & Salzberg, S. L. TopHat: discovering splice junctions with RNA-Seq. Bioinformatics (Oxford, England) 25, 1105–1111, doi:10.1093/bioinformatics/btp120 (2009).

53 Trapnell, C. et al. Transcript assembly and quantification by RNA-Seq reveals unannotated transcripts and isoform switching during cell differentiation. Nature biotechnology 28, 511–515, doi:10.1038/nbt.1621 (2010).

54 Ramskold, D., Wang, E. T., Burge, C. B. & Sandberg, R. An abundance of ubiquitously expressed genes revealed by tissue transcriptome sequence data. PLoS computational biology 5, e1000598, doi:10.1371/journal.pcbi.1000598 (2009).

55 Corradin, O. et al. Combinatorial effects of multiple enhancer variants in linkage disequilibrium dictate levels of gene expression to confer susceptibility to common traits. Genome research 24, 1–13, doi:10.1101/gr.164079.113 (2014).

56 Reimand, J., Arak, T. & Vilo, J. g:Profiler--a web server for functional interpretation of gene lists (2011 update). Nucleic Acids Res. 39, W307–315, doi:10.1093/nar/gkr378 (2011).

57 Merico, D., Isserlin, R., Stueker, O., Emili, A. & Bader, G. D. Enrichment map: a network-based method for gene-set enrichment visualization and interpretation. PLoS One 5, e13984, doi:10.1371/journal.pone.0013984 (2010).

58 Song, L. & Crawford, G. E. DNase-seq: a high-resolution technique for mapping active gene regulatory elements across the genome from mammalian cells. Cold Spring Harbor protocols 2010, pdb.prot5384, doi:10.1101/pdb.prot5384 (2010).

59 van de Werken, H. J. et al. Robust 4C-seq data analysis to screen for regulatory DNA interactions. Nature methods 9, 969–972, doi:10.1038/nmeth.2173 (2012).

60 Liu, T. et al. Cistrome: an integrative platform for transcriptional regulation studies. Genome biology 12, R83, doi:10.1186/gb-2011-12-8-r83 (2011).

61 Knott, S. R. et al. A computational algorithm to predict shRNA potency. Molecular cell 56, 796–807, doi:10.1016/j.molcel.2014.10.025 (2014).

62 Zuber, J. et al. Toolkit for evaluating genes required for proliferation and survival using tetracycline-regulated RNAi. Nature biotechnology 29, 79–83, doi:10.1038/nbt.1720 (2011).

63 Goecks, J., Nekrutenko, A. & Taylor, J. Galaxy: a comprehensive approach for supporting accessible, reproducible, and transparent computational research in the life sciences. Genome biology 11, R86, doi:10.1186/gb-2010-11-8-r86 (2010).

64 Osborne, T. S. et al. Evaluation of eIF4E expression in an osteosarcoma-specific tissue microarray. Journal of pediatric hematology/oncology 33, 524–528, doi:10.1097/MPH.0b013e318223d0c1 (2011).

65 Sakuma, T. et al. Efficient TALEN construction and evaluation methods for human cell and animal applications. Genes Cells 18, 315–326, doi:10.1111/gtc.12037 (2013).

66 Cermak, T. et al. Efficient design and assembly of custom TALEN and other TAL effector-based constructs for DNA targeting. Nucleic Acids Res. 39, e82, doi:10.1093/nar/gkr218 (2011).

